# RAREsim: A simulation method for very rare genetic variants

**DOI:** 10.1101/2021.04.13.439644

**Authors:** Megan Null, Josée Dupuis, Christopher R. Gignoux, Audrey E. Hendricks

## Abstract

Identification of rare variant associations is crucial to fully characterize the genetic architecture of complex traits and diseases. Essential in this process is the evaluation of novel methods in simulated data that mirrors the distribution of rare variants and haplotype structure in real data. Additionally, importing real variant annotation enables in silico comparison of methods that focus on putative causal variants, such as rare variant association tests, and polygenic scoring methods. Existing simulation methods are either unable to employ real variant annotation or severely under- or over-estimate the number of singletons and doubletons reducing the ability to generalize simulation results to real studies. We present RAREsim, a flexible and accurate rare variant simulation algorithm. Using parameters and haplotypes derived from real sequencing data, RAREsim efficiently simulates the expected variant distribution and enables real variant annotations. We highlight RAREsim’s utility across various genetic regions, sample sizes, ancestries, and variant classes.

## Introduction

Studies of rare variants are important to gain a full understanding of the genetics of health and disease, informing targeted drug development and precision medicine. Rare variants (minor allele frequency (MAF) <1%) have been associated with traits across many diseases including cancer, kidney, neurodevelopmental, cardiovascular, and infectious disease. With decreasing sequencing costs, rare variant data are increasingly accessible^1^ resulting in large sequencing studies (e.g. >35,000; 45,000; and 70,000 subjects), databases including the UKBiobank, GenomeAsia, and NIH programs such as the Genome Sequencing Program (GSP) and Trans-Omics for Precision Medicine (TOPMed). Rare variant methods continue to be developed (e.g. SKAT-O, iECAT, ProxECAT, and ACAT) to take advantage of the ever increasing sequencing data.

Simulation studies enable evaluation of methods and study design (e.g. power and sample size estimates) in known and controlled settings. Simulations that do not adequately mirror essential properties of real data may have issues generalizing to real data, potentially resulting in incorrect conclusions of method efficacy or power. In general, four qualities are necessary to emulate in rare variant simulations of a genetic region: (1) allele frequency spectrum (AFS), (2) total number of variants, (3) haplotype structure, and (4) variant annotation. To our knowledge, no rare variant simulation method currently exists that incorporates all four qualities.

1. **AFS** is the distribution of variant allele frequencies within a genetic region. Numerous studies have shown that the AFS is skewed towards very rare variants with the vast majority of variants being singletons and doubletons^2-4^.
2. The **total number of variants**, especially very rare variants, differs by ancestry and sample size. The total number of known variants is expected to increase as more ancestrally diverse and larger samples are sequenced^5,6^.
3. **Haplotype structure**, the linkage disequilibrium (LD) and probability that rare single nucleotide variants (SNVs) appear on the same haplotype background, varies across the genome and by ancestry.
4. **Variant annotation** is often used in rare variant methods and is thus essential for accurate evaluation of those methods. For instance, weighting functional variants has been shown to increase power to detect rare variant association in a gene region^1,7,8^. Other variant annotation, such as association with disease, is used to evaluate pleiotropy and in genetic correlation and polygenic risk scores. A great variety of variant annotation exists such as functional consequences, conservation score, chromatin state, eQTLs, epigenetic information, and prior disease associations, among others^9^. While *in silico* simulation of variant annotation can capture and emulate some annotation patterns, simulations derived from real data can easily incorporate precise empirical patterns from multiple annotation types, even those unique to a specific genetic region of interest, providing a more direct link between simulations and real data.

Population genetics simulation methods, such as Wright-Fisher^10,11^ and coalescent^12^, require only demographic and recombination information as input and often achieve an AFS, total number of variants, and LD structure similar to real data. However, these methods can be extremely computationally expensive or are not designed to emulate existing genetic regions resulting in an inability to use real variant annotations. Alternatively, resampling methods create haplotype mosaics from real genetic data using techniques that mimic recombination and mutations, maintaining the ability to use existing annotations. These methods, such as HAPGEN2^13^ derived from the original work of Li and Stephens^14^, are relatively computationally efficient and maintain the appropriate AFS, expected number of variants, and haplotype structure when simulating common variants^13,15^. However, as we and others^16^ show, HAPGEN2 does not simulate the correct total number or AFS for rare variants simulating too few rare and very rare variants (e.g. singletons and doubletons). There is currently no available software to simulate rare variant genetic data with a realistic AFS while retaining variant annotation.

To address this gap, we present RAREsim, a flexible and scalable genetic simulation method designed for accurate simulation of rare variants. We assess and show the utility of RAREsim across a variety of genetic regions, datasets, ancestries, and sample sizes. We provide RAREsim as an R package to enable easy implementation and appropriate simulation of rare variant data.

## Results

### Algorithm

RAREsim uses two primary datasets: input simulation data and target data. The *input simulation dataset* is a sample of haplotypes (e.g. 1000 Genomes haplotypes^3^) with minor alleles coded as 1 and all reference alleles, including monomorphic bases, coded as 0. The *target dataset* is summary level data used to estimate RAREsim parameters. The target data has two components: the allele count at each variant and the total number of variants in a genetic region of interest at various sample sizes (e.g. downsamplings from gnomAD^2^). While the *input simulation dataset* is required, the *target dataset* is not necessary if default or user defined parameters are used.

RAREsim has three main steps: (1) simulating haplotypes, (2) estimating expected number of variants, and (3) pruning rare variants to match expected. A flowchart summarizing the RAREsim algorithm is in **Figure 1**.

**Figure 1.**
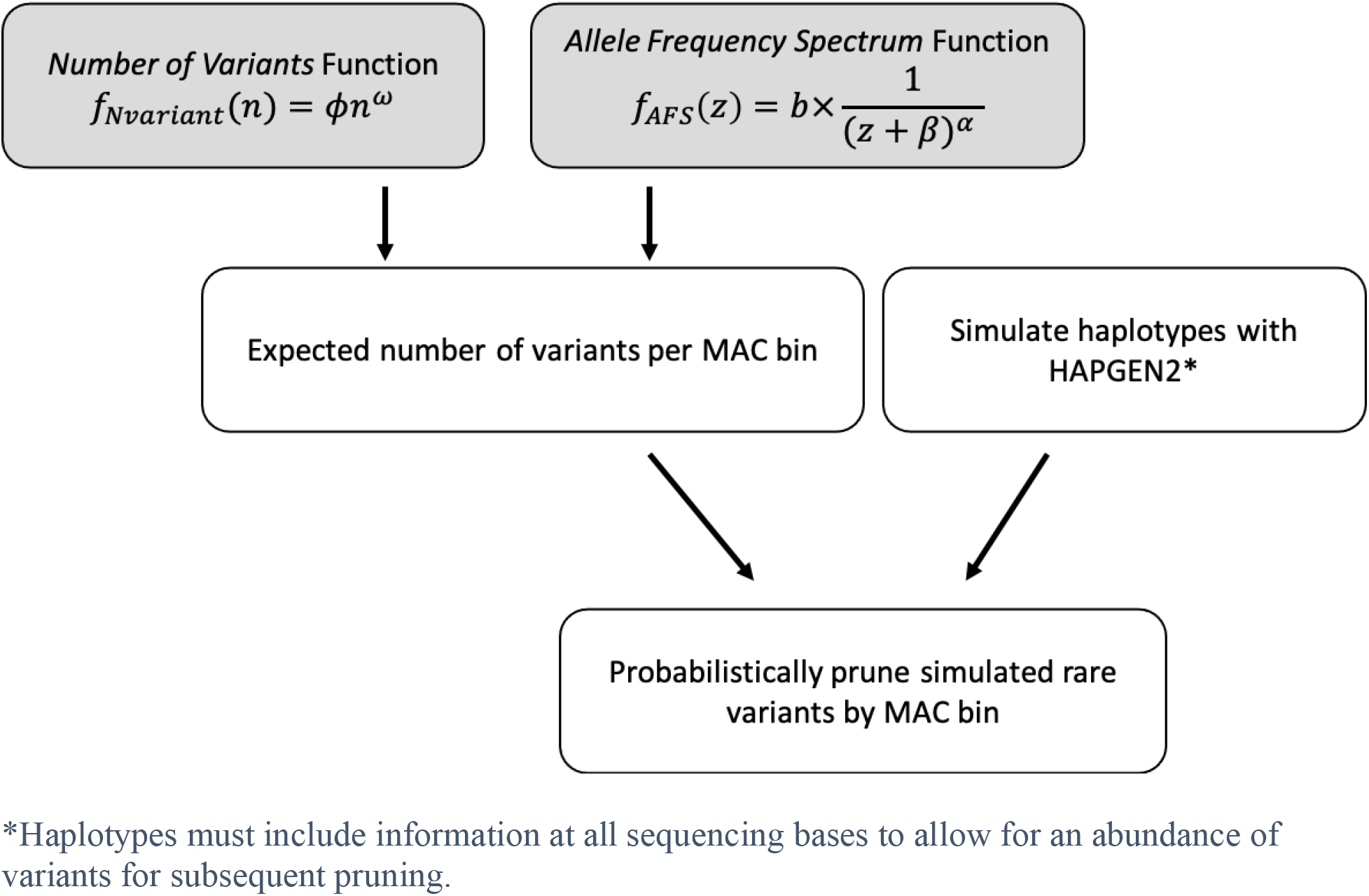
Flowchart of RAREsim. Flowchart describing RAREsim simulation process. Simulation parameters can be estimated using target data, default parameters, or user defined parameters (gray).

#### (1) Simulating an abundance of rare variants

RAREsim uses HAPGEN2^13^ to simulate haplotypes for *N_sim_* individuals. HAPGEN2 simulates haplotypes by creating mosaics of input and already simulated haplotypes, using recombination information so that regional LD is retained^13,15^. When all sequencing bases, including monomorphic, are included in the input haplotypes more de novo variants are simulated than expected. By sampling from previously simulated haplotypes to create a new haplotype, HAPGEN2 resamples de novo variants resulting in inflated numbers of rare variants.

#### (2) Estimating expected number of variants per MAC bin

The number of variants per MAC bin is estimated using two functions. The parameters for these functions are estimated using target data (described below). Alternatively, user-defined or default parameters can be used, eliminating the need for the user to provide and fit target data. The default parameters were derived using the default target data **(Methods)**.

##### (2a) Number of Variants Function

The total number of variants in a region depends on the sample size. RAREsim estimates the expected number of variants per kilobase (Kb) for a sample size *n* using the *Number of Variants* function. Specifically,

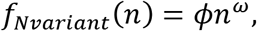

where *f_Nvariant_*(*n*) is the number of variants per Kb for *n* individuals. The parameters *ϕ* and *ω* are estimated to modify the scale and shape of the function, respectively. When simulating *N_sim_* individuals, RAREsim calculates the total number of variants in the region by multiplying the size of the region in Kb, *S_Kb_*, by the expected number of variants per Kb, *f_Nvariant_*(*n* = *N_sim_*).

The target data used for the *Number of Variants* function provides the observed number of variants per Kb, *T_n_*, in the simulation region observed at sample size *n*. The parameters are optimized by minimizing a least squares loss function summed over all observed sample sizes in the target data:

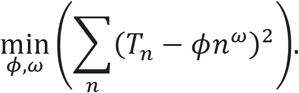

Sequential quadratic programming (SQP) via the *slsqp* function in the nloptr R package^17^ is used with constraints 0 < *ω* < 1 and *ϕ* > 0 to minimize the loss function. The initial starting values for the algorithm are *ω* = 0.45 and *ϕ* such that the largest observed sample size, *n_max_*, fits the observed number of variants per Kb at *n_max_*, 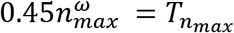. If the initial starting values do not result in a sufficient fit (loss > 1,000), a range of starting values are evaluated: *ω* ∈ {0.15,0.25,0.35,0.45,0.55,0.65}.

##### (2b) Allele Frequency Spectrum Function (AFS Function)

The *AFS* function, *f_AFS_*(*z*), estimates the proportion of variants with MAC = *z*. The largest MAC, *z_max_*, has MAF ≈1% in the target dataset. For a target dataset with *N_target_* individuals, *z_max_* = *floor*(*N_target_* * 2 * 0.01). Specifically,

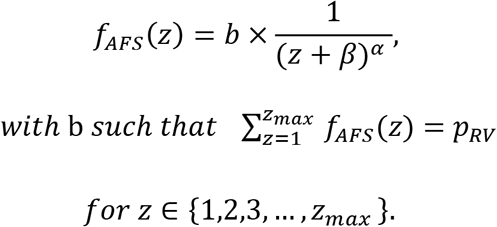

The scale parameter *b* ensures the sum of all individual rare MAC proportions equals the total proportion of rare variants observed in the target data, *p_RV_*. Parameters *α* and *β* determine the shape of the distribution.

Because individual MAC = *z* may have no observed minor alleles in the target data, particularly for higher rare MACs (e.g. MAC=10, 11, 12), MAC bins are used to optimize parameters. MAC bins are mutually exclusive, exhaustive groups of rare MACs. Seven MAC bins are used here: singletons, doubletons, MAC = 3-5, MAC = 6-10, MAC = 11-20, MAC = 21-MAF = 0.5%, MAF = 0.5% - 1%, denoted as *Bin*_1_,*Bin*_2_,…,*Bin*_7_, respectively. The total number and thresholds to define the bins can be modified by the user. Within each bin *j*, the estimated proportion of variants, 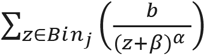, is compared to the observed proportion of variants in the target data, ∑_*z*∈*Bin_j_*_ *A_z_*. *A_z_* is the observed proportion of variants with MAC = *z*. Parameter estimates for *α*. and *β* are found by minimizing the least squares loss over all bins using SQP^17^ constraining *α* > 0,

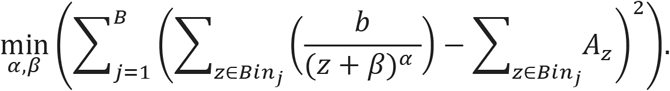

##### (2c) Expected Number of Variants per Minor Allele Count Bin

RAREsim uses the total number of variants within a genetic region for *N_sim_* individuals, *f_Nvariant_*(*N_sim_*) * *S_Kb_*, and the proportion of variants in *Bin_j_* to obtain the expected number variants in each MAC bin, 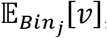,

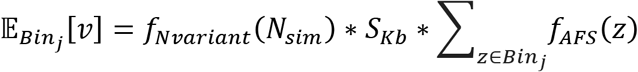

The expected total number of rare variants is calculated by summing across all rare MAC bins.

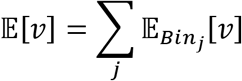

The total number of simulated rare variants (*M_sim_*) is calculated by summing over all MAC bins.

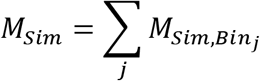

#### (3) Pruning

As described in (1), simulations using HAPGEN2 usually result in a larger total number of simulated rare variants than expected from step **(2)**. Simulated variants are pruned by retuning all or a subset of alternate alleles to reference alleles. Within HAPGEN2 and similar to real haplotypes, rare alleles have a high probability of being on the same haplotype background. Pruning alternate alleles preserves the high likelihood that rare alleles are on the same haplotype. Variants are probabilistically pruned, creating variability over the simulation replicates in the number of variants per MAC bin.

RAREsim sequentially prunes variants from high to low MAC bins starting with the highest rare MAC bin that has at least 10% more simulated variants than expected (i.e. 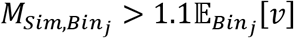).

##### (1) For MAC bins with more simulated variants than expected

a simulated variant is pruned with probability *P*(*rem*)_*j*_, where

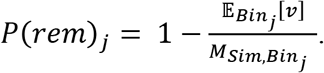

For each variant within the bin, RAREsim randomly draws from a Uniform(0,1) distribution. If the draw is within [0, *P*(*rem*)_*j*_], the variant is pruned. The location of a pruned variant is stored to allow variants to be added back at lower MAC bins that have fewer simulated variants than expected, as described in the next section.

##### (2) For MAC bins with fewer simulated variants than expected

(i.e. 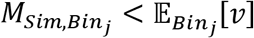), each of the *K* previously pruned variants from higher MAC bins are added to *Bin_j_* with probability,

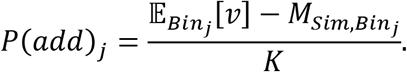

Random draws from Uniform(0,1) are used to determine which variants to add. The variant is added if the draw is within [0, *P*(*add*)_*j*_]. The MAC for each added variant is determined with a random sample from all possible MACs within *Bin_j_*. RAREsim then randomly samples without replacement the necessary number of haplotypes containing the alternate allele for the given variant. The allele for all other haplotypes is returned to reference.

## Results

### Evaluation of Number of Variants and Allele Frequency Spectrum Functions

*AFS* and *Number of Variants* functions were fit using target data from gnomAD v2.1 for four ancestry/sample-size groups (African, N = 8,128; East Asian, N = 9,197; Non-Finnish European, N=56,885; South Asian, N = 15,308) (**Supplemental Table 1**). Data from chromosome 19 was divided into 1 cM blocks; six blocks were merged with the preceding adjacent block (**Methods**), resulting in 101 blocks for simulation (**Supplemental Table 2**).

The ancestry/sample-size specific fitted *Number of Variants* function closely matches the observed values for all four ancestries (**Figure 2**, **Supplemental Figure 1**). Ninety percent of cM blocks had a relative difference within 2.42% for the estimated vs. observed number of variants per Kb. The average relative difference for all ancestries was *-1.10%* (90% CI = (−1.17%, −1.02%)), with ancestry specific averages of −0.65% (African, 90% CI = (−0.83%, −0.47%)), −1.09% (East Asian, 90% CI = (−1.24%, −0.94%)), −1.64% (Non-Finnish European, 90% CI = (−1.74%, −1.55%)), and −1.00% (South Asian, 90% CI = (−1.13%, −0.87%)) (**Supplemental Figure 2**). The negative mean relative differences indicate a slight but systematic overestimation of the number of variants per Kb for most blocks. Within a given target dataset, the *Number of Variants* function appears to slightly overestimate the observations of larger sample sizes and underestimate the observations of smaller sample sizes (**Supplemental Figure 1**).

**Figure 2.**
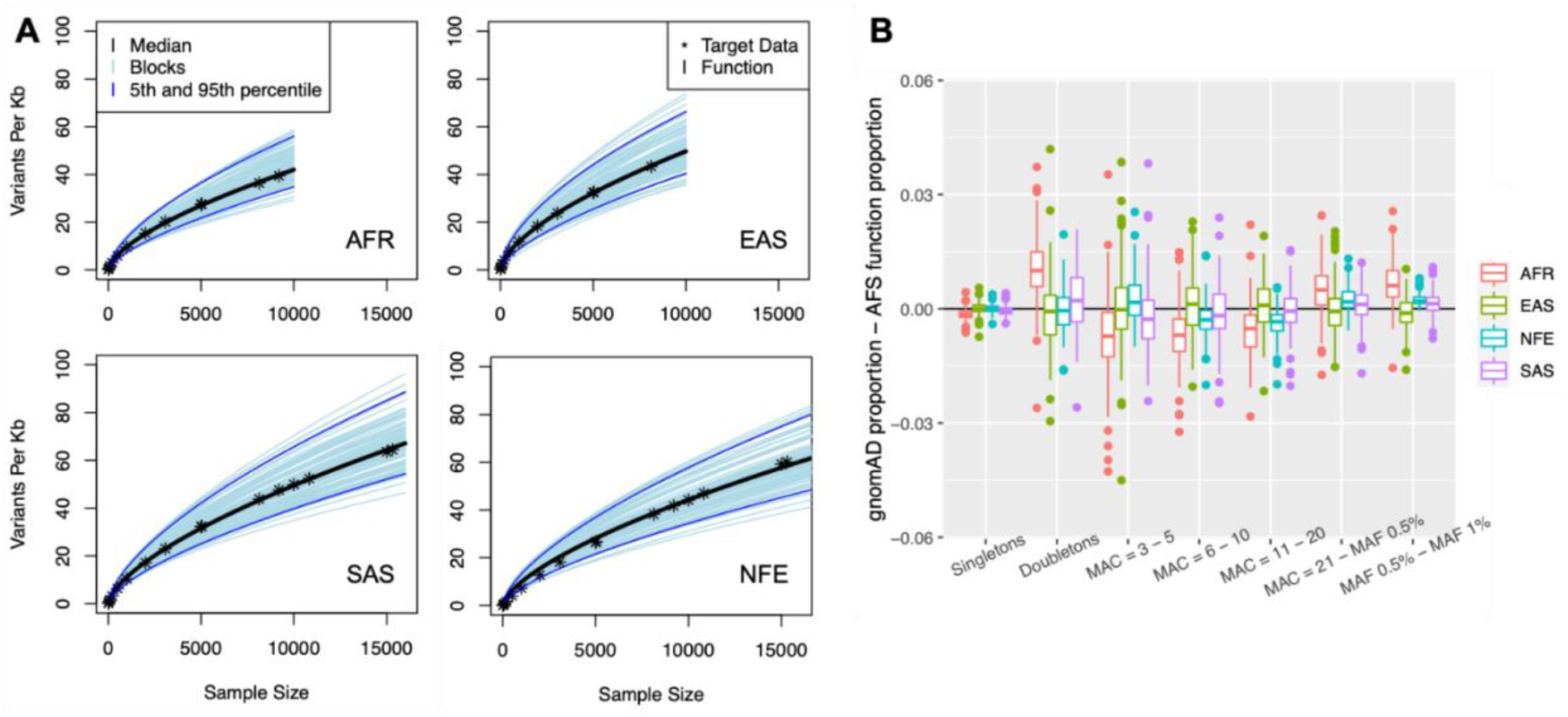
Evaluation of function fit. **A) *Number of Variants* function:** Fitted *Number of Variants* functions for all cM blocks. The median block is shown in black and the 5^th^ and 95^th^ blocks are shown in dark blue. The observed target data (*) for the median block are close to the fitted function for all four ancestries. Sample sizes up to 15,000 are shown here. The full sample size for the Non-Finnish European sample is in Supplemental Figure 1. **B) *AFS* function:** Difference between the gnomAD target data and estimates from the *AFS* function for the proportion of variants in each MAC bin for all chromosome 19 blocks by ancestry and MAC bin. All absolute differences are within 0.05, indicating the *AFS* function fits the target data well.

Observed and estimated variation in the number of variants per Kb across cM blocks increases with sample size (**Figure 2A**). However, even for the largest available target data sample size (Non-Finnish European, N=56,885), the variability of the number of variants per Kb remains low with 90% of the block-specific estimates within 35 variants of the median estimate.

The *AFS* function matched the observed data well with no apparent systematic bias. The average absolute difference between the observed and estimated proportion of variants in each MAC bin over all ancestries, blocks, and MAC bins was 0.53% (90% CI = (0.51%, 0.55%)) (**Figure 2B**). Ninety percent of the estimated proportions were within 1.3% of that observed. The maximum absolute difference in MAC bin proportion was 4.50% (observed in East Asian, MAC 3-5). Singleton counts matched particularly well, with a maximum absolute difference of 0.73%.

Despite different ancestries and widely different sample sizes (from N = 8,128 to N = 56,885 for African and Non-Finnish European respectively) the proportion of variants per MAC bin were similar (**Supplemental Figure 3**). There is more variation between ancestry/sample-size groups for the total proportion of rare variants (i.e. proportion of all variants with MAF <1%) (**Supplemental Figure 4**). Regardless, within each ancestry, variation of the AFS between cM blocks remains small. For instance, the MAC bin with the most variation (East Asian singleton bin) has 90% of the blocks having estimated proportions within 6.3% of the median.

### Evaluation of Simulation Results

One hundred replicates of each block were simulated using RAREsim and HAPGEN2 matching the gnomAD sample size for each ancestry group (*N_African_* = 8,128, *N_East Asian_* = 9,197, *N_Non–finnish European_* 56,885, *A_South Asian_* 15,308) (**Figure 3**). RAREsim produced a similar number of variants relative to gnomAD across all ancestry groups and MAC bins indicating that the total number of variants and AFS are representative of real sequencing data. Conversely, HAPGEN2^13^ with only polymorphic SNVs greatly underestimated the total number of rare variants, especially very rare variants. HAPGEN2 simulations including all sequencing bases produced many more rare variants than observed. These results are consistent across all cM blocks and for the cumulative chromosome 19 coding region (**Supplemental Figures 5-8, Supplemental Table 3-6**).

**Figure 3.**
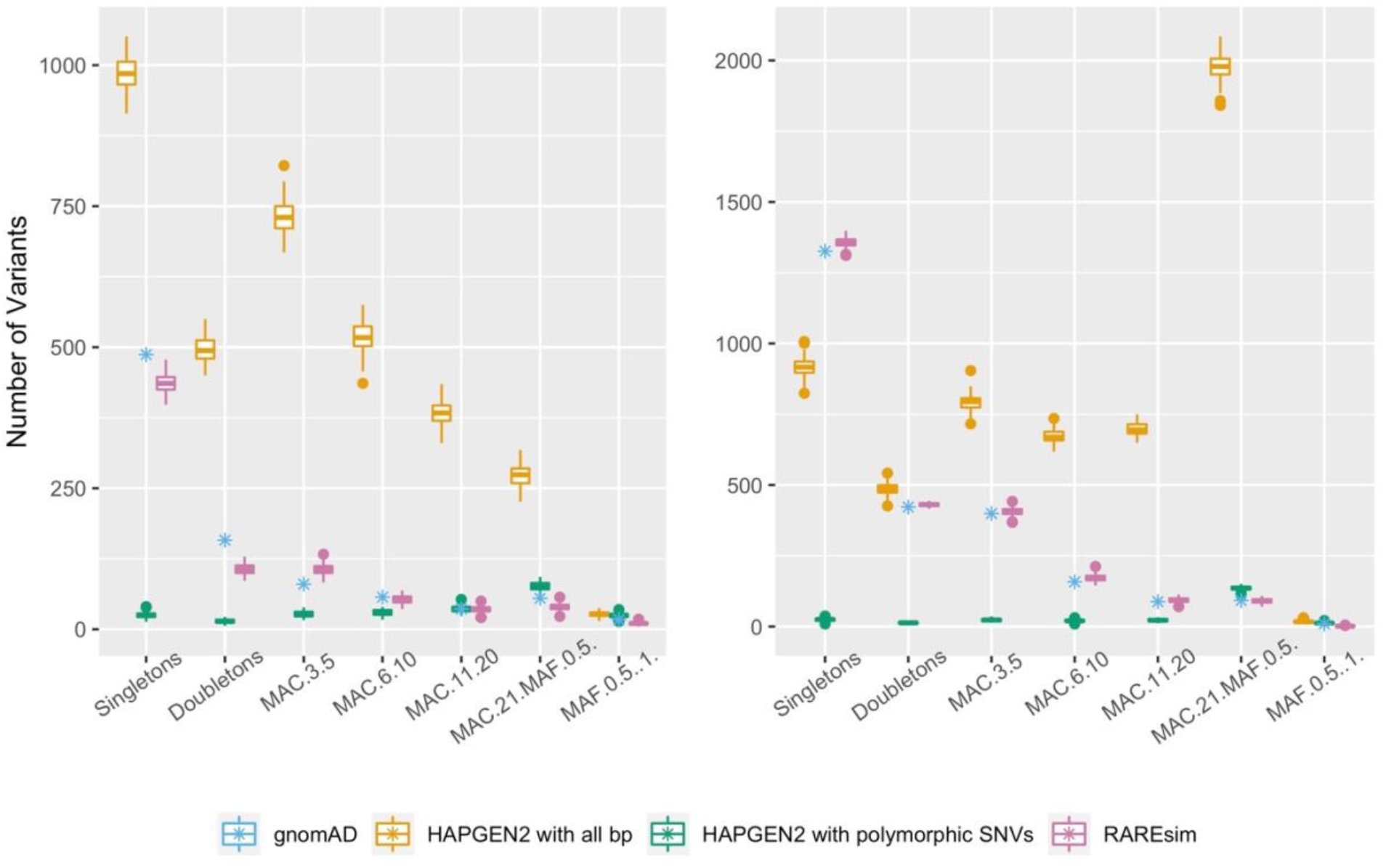
Evaluation of RAREsim. The distribution and number of variants simulated using RAREsim (pink), HAPGEN2 with only polymorphic SNVs (default, green), and HAPGEN2 with all sequencing bases (yellow) is compared to gnomAD (blue) for the cM block with the median number of bp. Ancestry specific simulations are shown for African (N = 8,128; left) and Non-Finnish European (N = 56,885; right) matching the sample size observed in gnomAD v2.1. RAREsim emulates the expected number of variants within each MAC bin, while the other simulation methods either grossly underestimate (HAPGEN2 with polymorphic SNVs) or overestimate (HAPGEN2 with all sequencing bp) the number of variants.

### Generalizability of Default Parameters

Ancestry/sample-size specific default parameters for the *Number of Variants* and *AFS* functions (**Table 1**) were estimated using the median observation over cM blocks for each ancestry/sample-size group. RAREsim default parameters performed well, matching the observed number of variants and AFS in a wide variety of situations including three GENCODE regions on chromosomes 1, 6 and 9^18^, in non-coding regions within blocks, and in other datasets/sample sizes - gnomAD v3 and UKBioBank^19^ (**Methods, Figure 4, Supplemental Figures 9 – 14**). The simulated sample sizes evaluated were up to ~3x larger (gnomAD v3African) and ~2x smaller (gnomAD v3 Non-Finnish European) than the sample sizes used to derive the default parameters. The default parameters often performed similarly to the cM block specific simulation parameters and always outperformed HAPGEN2.

**Table 1:** Default function parameters estimates.

| | Number of Variants<br>$f_{Nvariant}(x) = \phi x^{\omega}$ | | Allele Frequency Spectrum<br>$f_{AFS}(z) = b \times \frac{1}{(z + \beta)^{\alpha}}$ | | |
| --- | --- | --- | --- | --- | --- |
| | $\hat{\phi}$ | $\hat{\omega}$ | $\hat{\alpha}$ | $\hat{\beta}$ | $\hat{b}$ |
| **African** | 0.1576 | 0.6247 | 1.5883 | -0.3083 | 0.2872 |
| **East Asian** | 0.1191 | 0.6369 | 1.6656 | -0.2951 | 0.3137 |
| **Non-Finnish European** | 0.1073 | 0.6539 | 1.9470 | 0.1180 | 0.6676 |
| **South Asian** | 0.1249 | 0.6495 | 1.6977 | -0.2273 | 0.3564 |

**Figure 4.**
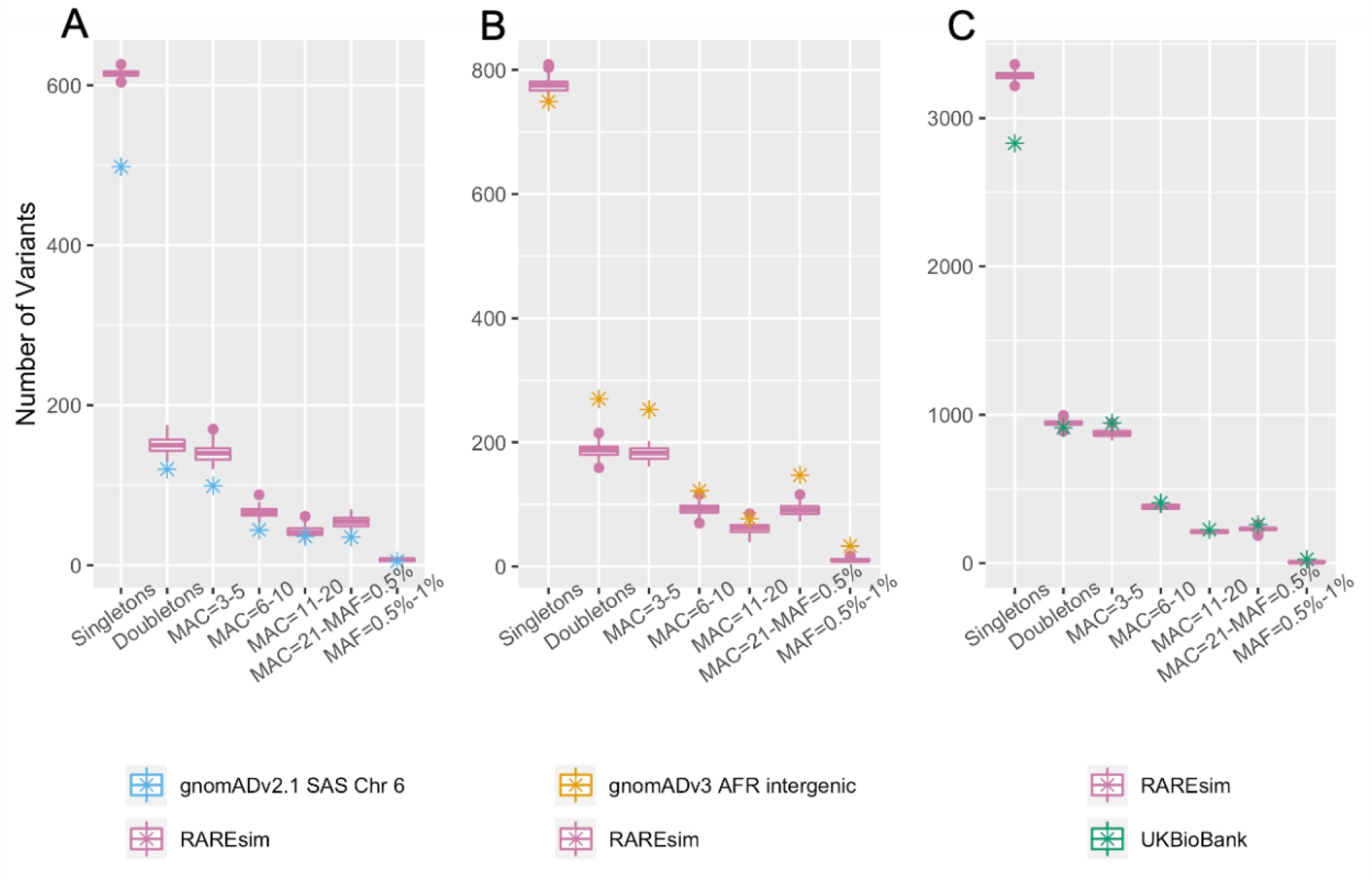
Generalizability of Default Parameters. Utility of RAREsim’s ancestry specific default parameters for different chromosomes (A), sample sizes (B & C), intergenic regions (B), and other target datasets (B & C). RAREsim simulations closely approximate the observed number of variants (y-axis) in each MAC bin (x-axis) in all scenarios. **A)** Simulations using South Asian default parameters for Chromosome 6 GENCODE region. **B)** Simulating a sample size of 21,042 using African ancestry default parameters (derived from N=8,128) for in an intergenic region from gnomAD v3. **C)** Simulating a sample size of 41,246 to match a British sample from the UK Biobank using Non-Finnish European default parameters (derived from N=56,885).

### Stratified Simulation of Functional and Synonymous Variants

As expected, we observed more functional SNVs than synonymous^2^, with the largest differences observed at MAC ≤ 5 (**Supplemental Figure 15**). This resulted in substantially different fitted *Number of Variants* functions for the two types of variants (**Supplemental Figure 16**). Stratified simulation of functional and synonymous variants closely approximated the number of variants observed in each MAC bin and suggests utility in separately simulating different groups of variants (**Supplemental Figure 17**).

### Simulation of Large Sample Sizes

As discussed previously (*Generalizability of Default Parameters*), RAREsim accurately simulated 21,042 African samples to match gnomAD v3 using ancestry specific default parameters derived from African gnomAD v2.1 (N=8,128). We currently assume that the AFS does not change with sample size. Consistent shape of AFS was observed over the gnomAD v2.1 ancestry/sample size groups (*N_African_* = 8,128, *N_East Asian_* 9,197, *N_Non–Finnish European_* 56,885, *N_South Asian_* = 15,308). Further, ancestry specific fitted *Number of Variants* functions extrapolated to larger sample sizes than observed were similar to the shape of the fitted *Number of Variants* function for the total gnomAD v2.1 sample (N=125,748) (**Supplemental Figure 18**). Therefore, we believe simulating sample sizes up to ~125,000 is likely reasonable.

### Computation Time

The time to simulate one replicate using a desktop with 32GB RAM for a cM block on chromosome 19 varied between 15 seconds for the smallest block (3,183 bp) and sample size (N_African_=8,128) and 12 hours 32 minutes 20 seconds for the largest block (81,235 bp) and sample size (N_Non-Finnish European_=56,5885). The median run time was 4 minutes 16 seconds (**Supplemental Table 7**).

The amount of time to simulate haplotypes with RAREsim is dependent on the number of samples being simulated and the size of the region. Simulating a region with ~19 Kb varied between 1 minute 34 seconds for N=8,128 and 37 minutes 34 seconds for N=56,885. When simulating N=15,308 individuals, RAREsim simulations took between 16 seconds for a region of ~3 Kb and 11 minutes 31 seconds for a region of ~81 Kb. The rate limiting step in large simulations was HAPGEN2. For the largest region and sample size, HAPGEN2 took over 11 hours to simulate when using a machine with 32 GB RAM. The same region was simulated in ~1 hour and 19 minutes using 192 GB RAM indicating that memory capacity was reached using the original computing specs (**Methods**).

## Discussion

Here we present RAREsim, a rare variant simulation algorithm. Unlike HAPGEN2, which either severely under or over simulates the proportion of very rare variants, RAREsim simulates the expected proportion of rare and very rare variants across a variety of genetic regions, ancestries, and sample sizes. RAREsim produces simulations that match the expected AFS, total number of variants, and haplotype structure while enabling variant annotation. To our knowledge, no other existing simulation software is able to emulate real data in all of these areas. We show that RAREsim’s ancestry specific default parameters derived from the coding regions of chromosome 19 generalize to other chromosomes, datasets, sample sizes^19^, and non-coding regions, approximating the number of variants per MAC bin with remarkable accuracy. We offer user flexibility by enabling use of RAREsim with default parameters, user defined parameters, or parameters estimated to match user provided target data.

For typical uses of simulated genetic data (i.e. evaluating or comparing methods and general power analysis) we recommend simulating with the default parameters. Default parameters were shown to be robust across sample sizes, chromosomes, coding and intergenic regions, and datasets. It is possible, although we believe unlikely, that the default parameters will perform poorly when used in scenarios not evaluated here. If precise matching of a particular empirical data characteristic such as functional variant type, genetic region, sample size, or ancestry is important, we recommend re-estimating the simulation parameters using RAREsim functions. Additionally, parameters are able to be specified without fitting target data. For example, to simulate a specific ancestry, a user could make an educated decision on the total number of variants based on the relationship to the ancestries evaluated here (e.g. total number of variants between that of African and European).

RAREsim simulates haplotypes in the same form as HAPGEN2: hap/leg/sample files^20^. Haplotype files can be converted to vcf files using the SHAPEIT *convert* command^21^ or bcftools *haplegendsample2vcf* command^22^. Genetic association with disease can be simulated from a sample of generated haplotypes using an existing software such as PhenotypeSimulator^23^. Simulation of families or large pedigrees can be performed with a pedigree simulation software such as ped-sim^24^.

It has been shown that sample sizes in the tens to hundreds of thousands are needed to have sufficient power to detect associations with rare variants^25^. Due to the lack of very large (>100,000), ancestry specific, publicly available target data at the time of publication, we could not assess the accuracy of the RAREsim simulations for very large sample sizes. As genetic sequencing resources continue to increase in size, RAREsim will be ideally suited to simulate large sample sizes with estimation of new simulation parameters. For the *Number of Variants* function, the extrapolated, ancestry specific *Number of Variants* functions were compared with that of the full sample available in gnomAD v2.1. We believe that the *Number of Variants* function is able to accurately simulate sample sizes up to what is observed in gnomAD v2.1 (~125,000). Alternatively, users can use population-genetics theory or other resources to make an informed decision for the total number of variants expected for very large samples. One such resource is the Capture-Recapture^26^ algorithm, which can be used to estimate the number of segregating sites given allele count data. While Capture-Recapture can extrapolate to larger sample sizes, the software cannot be easily used for sample sizes that are smaller than the observed target data. RAREsim does not currently modify the *AFS* function as sample size increases. Consistent AFS were observed over the gnomAD sample sizes (N = 8,128 - 56,88). However, we expect the AFS to deviate with very large sample sizes. A user can update the *AFS* function parameters if desired, and research is ongoing to estimate the expected AFS for very large sample sizes. We believe that RAREsim can currently accurately simulate sample sizes up to ~125,000.

There are several limitations to RAREsim. First, RAREsim is only as good as the data on which the simulations are based. Errors or inconsistencies in the target data or input simulation haplotypes will be propagated through the simulations. Secondly, the default parameters were developed and evaluated on autosomes. A user may fit sex chromosome target data or assume parameters extend to sex chromosomes. Finally, RAREsim requires additional memory (e.g. >32 GB RAM) when simulating large regions and sample sizes. For efficient simulation of large regions and sample sizes, highmem computing or breaking up the simulation region into smaller portions and combining after simulation is needed. We are actively working on extensions for these limitations.

One of the primary benefits of RAREsim is its ability to match real target data, either provided by the user or as done here with gnomAD v2.1. Matching observed data allows RAREsim to adapt as sequencing data evolves due to technological advances or improved genetic resources from additional ancestral populations and increased sample sizes. For example, RAREsim will be able to approximate TopMED^27^ and ALFA, aggregated allele frequencies from dbGaP^28^ once these resources are released. RAREsim can also simulate unique characteristics of a specific genetic region such as haploinsufficiency, contribution to a polygenic risk score, or selection. The flexibility of RAREsim to emulate real data allows users to assess methods and complete power analyses for in relevant and realistic genetic regions and samples.

## Acknowledgements

The UK Biobank data was gathered using the UK Biobank Resource under Application Number 42614. We would like to thank Achilleas Pitsillides for obtaining the UK Biobank allele counts. We would also like to thank Robert Goedman for providing programming insight. We thank Dr. Ferdinand Baer for support of this project. This work was supported by the National Human Genome Research Institute (R35HG011293 and U01HG009080 to AEH and CGR; U01HG009080-05S1 to CGR).

## Methods

### Input Simulation Datasets

For the input simulation dataset, we used haplotypes, legend files (an accompanying variant list), and a recombination map from 1000 Genomes Phase 3 (hg19)^3^. The files were modified to include information at each sequencing base. The recombination map was derived from the combined sample of all ancestries (OMNI)^3^. Links to these resources are in **Supplemental Table 8**.

For the simulation haplotypes, African, East Asian, Non-Finnish European, and South Asian global ancestries were used with admixed African samples (African Caribbeans in Barbados (ACB) and Americans of African ancestry in Southwest Utah (ASW)) excluded. HAPGEN2 simulates biallelic SNVs. Hence, we omitted indels present in the 1000G haplotypes. For multiallelic SNVs, the first alternate allele in the legend file with at least one observed alternate allele was kept.

### Target Datasets

Exome sequencing data from gnomAD v2.1^2^ on chromosome 19 were used as target data to estimate the simulation parameters. For the *Number of Variants* function, the observed number of loss-of-function, synonymous, and missense variants by gene, ancestry group, and sample size (e.g. 500, 1000, 2000, 5000, etc.) from Karczewski et al.^2^ was used (https://storage.googleapis.com/gnomad-public/papers/2019-flagship-lof/v1.0/gnomad.v2.1.1.lof_metrics.downsamplings.txt.bgz, **Supplemental Table 8**). The number of variants for all three function classification groups were summed to obtain the total number of variants observed per gene. The GENCODE v19 file^18^ (link in **Supplemental Table 8**) contains the genomic positions of the canonical transcript coding regions used by Karczewski et al.^2^ to define genes. The total number of variants within the simulation region of interest was found by summing over all genes in the region. When a region contained overlapping genes, the proportion of overlap was calculated and removed from one gene, as to not count variants twice.

For the AFS target data, allele counts for biallelic SNVs in the coding region of canonical transcripts for four ancestry groups from gnomAD v2.1 were used (African, N = 8,128; East Asian, N = 9,197; Non-Finnish European, N=56,885; South Asian, N = 15,308)^2^ (https://storage.googleapis.com/gnomad-public/release/2.1.1/vcf/exomes/gnomad.exomes.r2.1.1.sites.vcf.bgz, **Supplemental Table 8**). We observed slight discrepancies for some regions in the total number of variants between the gnomAD v2.1 data used for the AFS target data and the gnomAD downsamplings data used for the Number of Variants target data (**Supplemental Figure 19**). Differences in the number of variants per gene likely arise due to inconsistencies between the two datasets with respect to classification of variant function, removal of variants in overlapping genes, as well as other differences. The discrepancies did not substantially affect the simulation results, as shown when the simulated haplotypes are compared to the gnomAD data (**Results**).

### Centimorgan Blocks

Chromosome 19 was divided into 1 cM blocks for simulation. The cM blocks were defined using the 1000 Genomes Project recombination map estimated from the combined set of all ancestries^3^ (*Input Simulation Datasets*). Blocks were restricted to the coding region of the canonical transcript for each gene. Genes that overlapped multiple blocks or were between cM blocks (the recombination map did not contain information at all bp) were included with the previous cM block. Of the 107 cM blocks, two blocks did not meet the requirement of containing at least two genes (blocks 17 and 23). Additionally, there were four blocks (blocks 8, 50, 57, and 92) with fewer than 100 SNVs in at least one ancestry in the gnomAD target data. These six blocks were merged with the preceding adjacent block, resulting in 101 blocks for simulation (**Supplemental Table 2**). Blocks ranged from 3,183 bp to 81,253 bp (median = 19,029; Q1 = 11,037; Q3 = 27,204).

### Implementation of RAREsim

RAREsim was implemented for a genetic region of interest using the computing flowchart shown in **Supplemental Figure 20**. First, the input haplotype and legend files were modified to include all bp, including monomorphic bases. Then, haplotypes were simulated for each cM block using HAPGEN2^13^ with default parameters. The relative risk was set to 1.0, and hence no disease loci were simulated. The random seed in HAPGEN is set by time. Therefore, simulation replicates cannot be run in parallel across multiple cores starting at the same time. To avoid this, we simulated replicates for the same simulation scenario on the same computing core in series. Alternatively, the *pause* Bash command could be used when simulating haplotypes in parallel. Then, the expected number of variants per MAC bin was calculated using the *expected_variants* RAREsim function with either default simulation parameters or region-specific simulation parameters estimated using *fit_afs* and *fit_nvariants* functions. Finally, the simulated haplotypes were pruned using the *prune_variants* function to identify pruning locations and a supplemental Bash script to efficiently prune the identified locations.

### Evaluation of Allele Frequency Spectrum and Number of Variants Functions

In our application, parameters for the *Number of Variants* and *AFS* functions were estimated for each of the 101 blocks and four ancestry groups from gnomAD. To evaluate how well the *Number of Variants* function fit the target data, we calculated the relative difference for the observed target data at the sample size available in gnomAD, *T_N_gnomAD__*, to the *Number of Variants* function estimate, 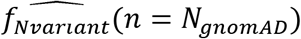. The relative difference was calculated as

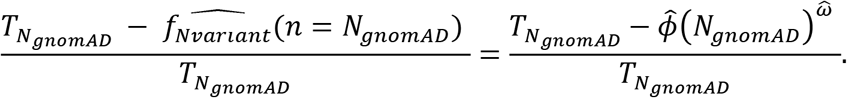

We evaluated fit of the *AFS* function with the difference between the estimated proportion of variants, 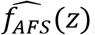, and the observed proportion in gnomAD, *A_z_*, for each MAC *Bin_j_*,

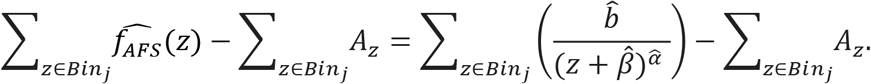

### Default Function Parameters

Ancestry specific default parameters were calculated using the median target data over all blocks (i.e. median number of variants per Kb at each sample size (*Number of Variants* function) and median proportion of variants in each MAC bin (*AFS* function)). For the *Number of Variants* function, the 5^th^ and 95^th^ percentile observations over the 101 blocks were used to estimate 5^th^ and 95^th^ percentile functions.

### Evaluation of Simulation Results

One hundred replicates of each block were simulated for the gnomAD sample size of each ancestry group (*N_African_* = 8,128, *N_East Asian_* 9,197, *N_Non–finnsh European_* 56,885, *N_South Asain_* = 15,308). The matched sample size enabled a direct comparison between gnomAD and simulated data as sample size greatly influences the number of variants expected in MAC bins. We compared RAREsim to the default implementation of HAPGEN2 with only polymorphic SNVs in the input simulation data and to HAPGEN2 using all bp, including monomorphic bp. Each block was simulated and pruned independently. To evaluate simulations from chromosome 19 as a whole, the variant counts for each MAC bin were summed over all cM blocks.

### Generalizability of Default Parameters

Generalizability of default parameters was assessed on different chromosomes, other sample sizes, in an intergenic region, and simulating data to match another dataset. To evaluate the performance of the default parameters for other chromosomes, we simulated GENCODE regions on chromosomes 1, 6, and 9 that were chosen to be representative of the genome^18^ (**Supplemental Table 9**). As with the blocks on chromosome 19, the regions were restricted to canonical coding exons. These blocks were each 500 Kb, but when restricted to the coding region were 24,918; 12,519; and 17,051 bp on chromosomes 1, 6, and 9 respectively.

Whole genome sequencing data from gnomAD v3 was used to evaluate the performance of default parameters for different sample sizes, and for intergenic regions. To evaluate default parameters for different sample sizes, we simulated three blocks (5^th^, 50^th^, and 95^th^ percentile block for number of variants) for the African ancestry group (N=21,042 for v3 compared to N=8,128 for v2.1) and Non-Finnish European ancestry group (N=32,299 for v3 compared to N=56,885 for v2.1). To evaluate the utility of default parameters for intergenic regions, we simulated intergenic regions within the three blocks limiting the original coding region size.

Finally, to evaluate performance of the default parameters in another dataset and sample size, we simulated a Non-Finnish European sample to match the UK Biobank^19^. Due to an error in the UK Biobank 50K release, the 95^th^ percentile block contained missing data; thus, we used the 5^th^, 50^th^, and 94^th^ percentile blocks instead. We simulated 41,246 Non-Finnish European individuals, which was the number of individuals in exome sequencing British sample after removing ethnic outliers.

### Stratified Simulation of Functional and Synonymous Variants

To demonstrate RAREsim’s ability to simulate different types of variants, such as variants in different functional classes, variants were stratified and simulated by functional and synonymous status. The reference and alternate allele are required for variant annotation. For polymorphic variants within gnomAD, the observed reference and alternate allele were used. Monomorphic bp in gnomAD were annotated using all possible alternate alleles with the *convert2annovar* function in ANNOVAR^29^. To restrict to one alternate allele, each allele was first annotated as a transition or transversion. Within the exome, Wang et al.^30^ observed transition to transversion ratios (Ti/Tv) between 2.79 and 2.84 across ancestries. Here, the average Ti/Tv of 2.815 was used to calculate the probability (0.7379) of a transition for each variant. For each monomorphic bp, we performed a random draw from a Uniform(0,1) distribution. If the random draw was within [0,0.7379], the transition alternate allele was used. Otherwise, the variant was annotated as a transversion and the alternate allele was assigned randomly from the two possible alternate alleles.

Variants were annotated using Ensembl Variant Effect Predictor (VEP)^31^ release 100. For variants with multiple annotations, the most severe consequence was chosen. Synonymous variants were those annotated as synonymous. Matching gnomAD’s annotation^2^, functional variants were those annotated as missense, frameshift, splice site disrupting, and stop gained. Stratified simulation with RAREsim by variant class, including refitting target data and pruning, was performed for the block with the median number of variants.

### Simulation of Large Sample Sizes

We fit the *Number of Variants* function to the median cM block for the total gnomAD v2.1 sample (N = 125,748) and compared the fitted, ancestry specific *Number of Variants* functions extrapolated to large sample sizes.

### Computing Time

Computing time was evaluated on an Ubuntu 18.04.2 LTS desktop with Intel^®^ Core TM i7-6700 CPG at CPU 8 x 3.40Ghz. The desktop is 64-bit with 1.1 TB (disk) GNOME 3.28.2 and 32GB RAM. For each ancestry, the simulation time for the cM block with the minimum (3,183), median (19,029), and maximum (81,235) bp was recorded. To re-evaluate the simulation of haplotypes with HAPGEN2 using more memory, a Dual Intel Xeon E5-2670v2 (2.5 Ghz x 10 cores, each), 192GB PC3-12800R RAM (12×16GB sticks) was used.

## Data Availability

All data used are publicly available with links found in **Supplemental Table 8**. The reference haplotype and legend files with all monomorphic sequencing bases within the coding regions included are available at https://github.com/meganmichelle/RAREsim_Example.

## Code Availability

RAREsim is an open-source R package, and all code can be found at https://github.com/meganmichelle/RAREsim. A small example simulation with the necessary script is available at https://github.com/meganmichelle/RAREsim_Example. Code to complete the majority of the analyses included here can also found at https://github.com/meganmichelle/RAREsim_Example.

## Author Contributions

M.N. and A.E.H. developed the algorithm and evaluation of the algorithm with insight from J.D. and C.R.G. All analyses were performed by M.N., and the R package was developed by M.N. M.N. and A.E.H. drafted the paper. A.E.H. supervised the project. All authors reviewed the final manuscript.

