## Supplemental Figures for "RAREsim: A simulation method for very rare genetic variants"

**RAREsim Supplemental Figures**

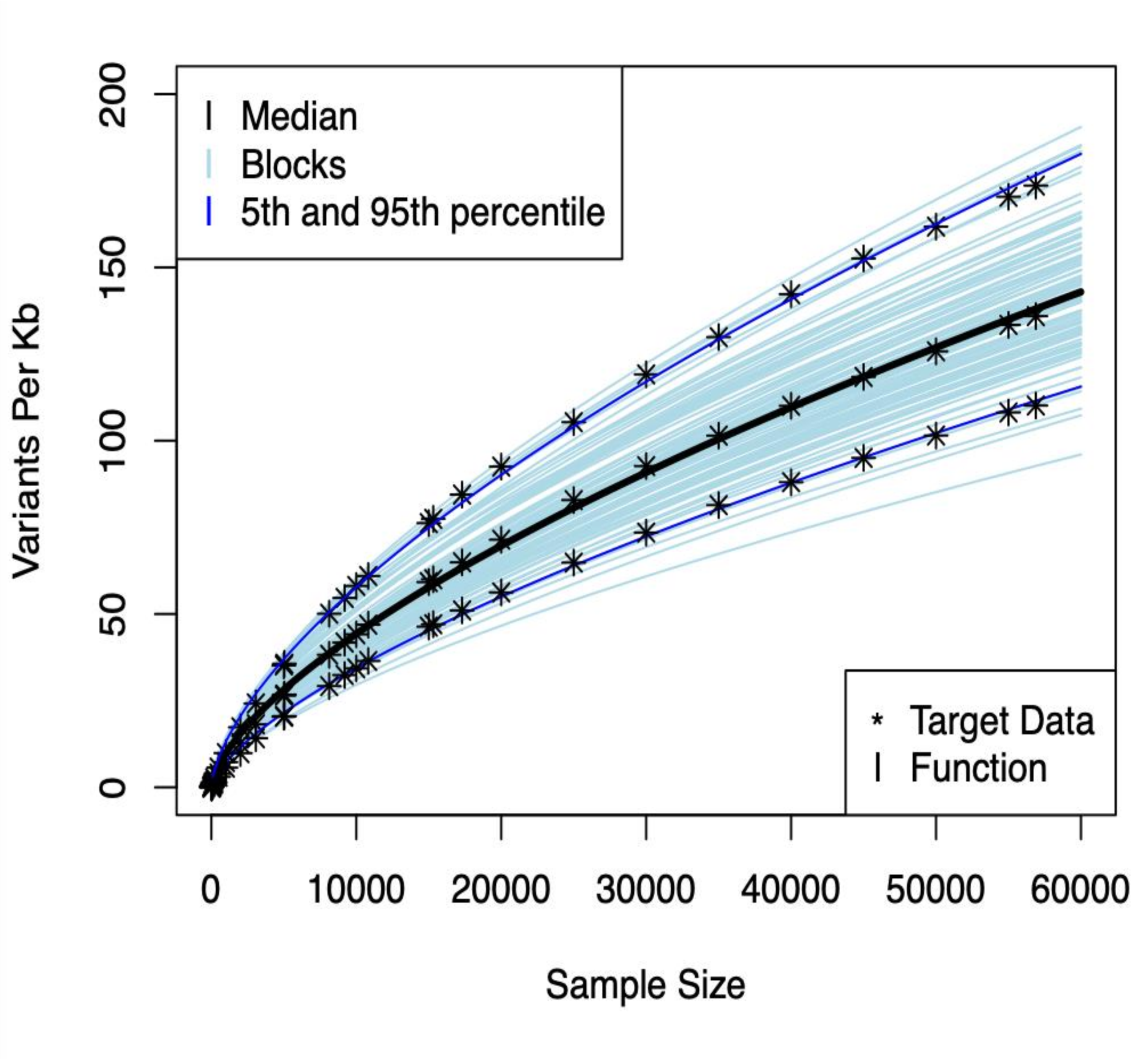


***Supplemental Figure 1. Number of Variants function for Non-Finnish European** The functions estimated from individual blocks are shown (light blue), with the median (black), and 5$^{th}$ and 95$^{th}$ percentile (dark blue) functions also shown. Observed target data are represented with stars for the median, 5$^{th}$, and 95$^{th}$ target data. The black fitted line – the default parameter function – and the dark blue lines fit their respective observed data very closely. We observe slight overestimation at the largest available sample size, with slight underestimation at smaller sample sizes.*

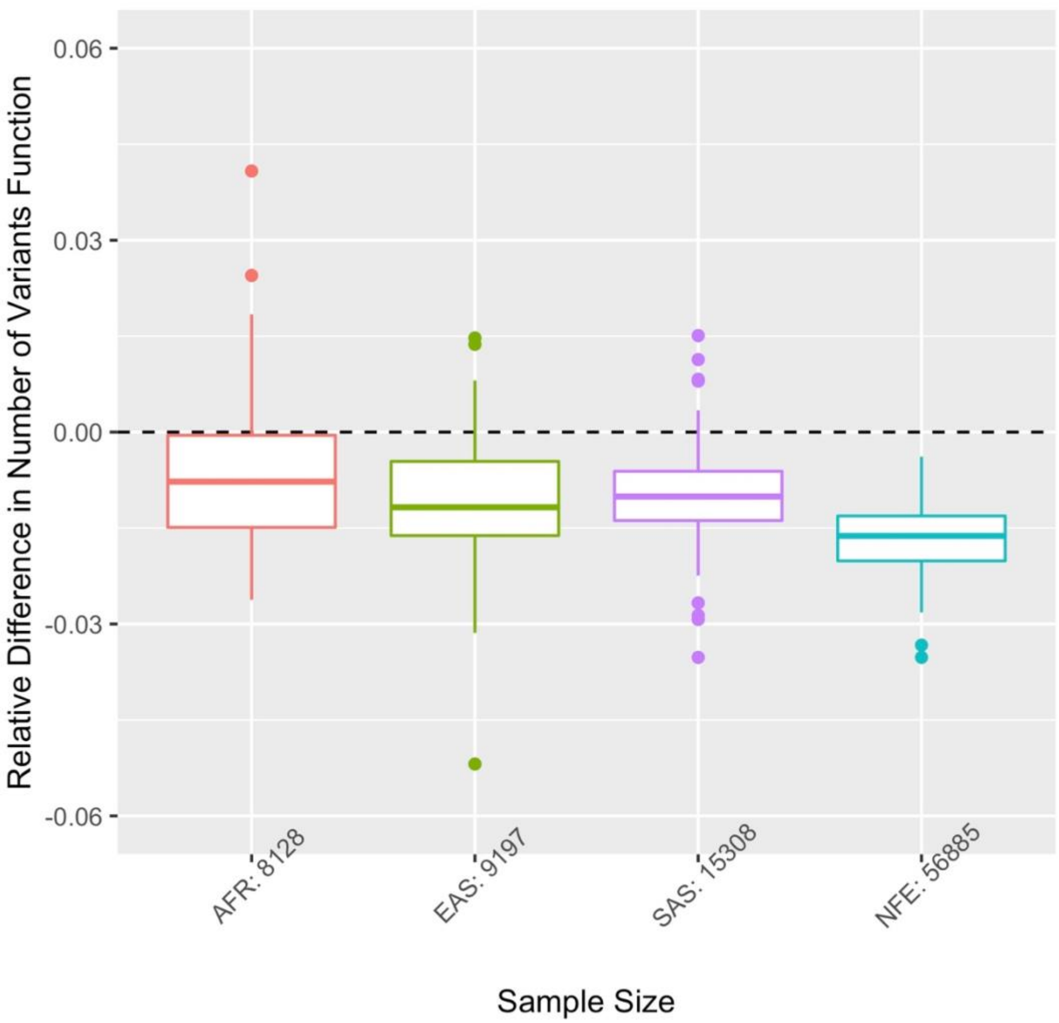


***Supplemental Figure 2. Relative difference of the Number of Variants function***
*The relative difference for a given sample size is the number of variants observed in gnomAD minus the function estimation, all divided by the number of variants observed in gnomAD. The Number of Variants function at the available sample size is gnomAD shows that there are systematically slightly less variants observed in gnomAD than the Number of Variants function estimates. The x-axis shows the ancestry and sample size available in gnomAD.*

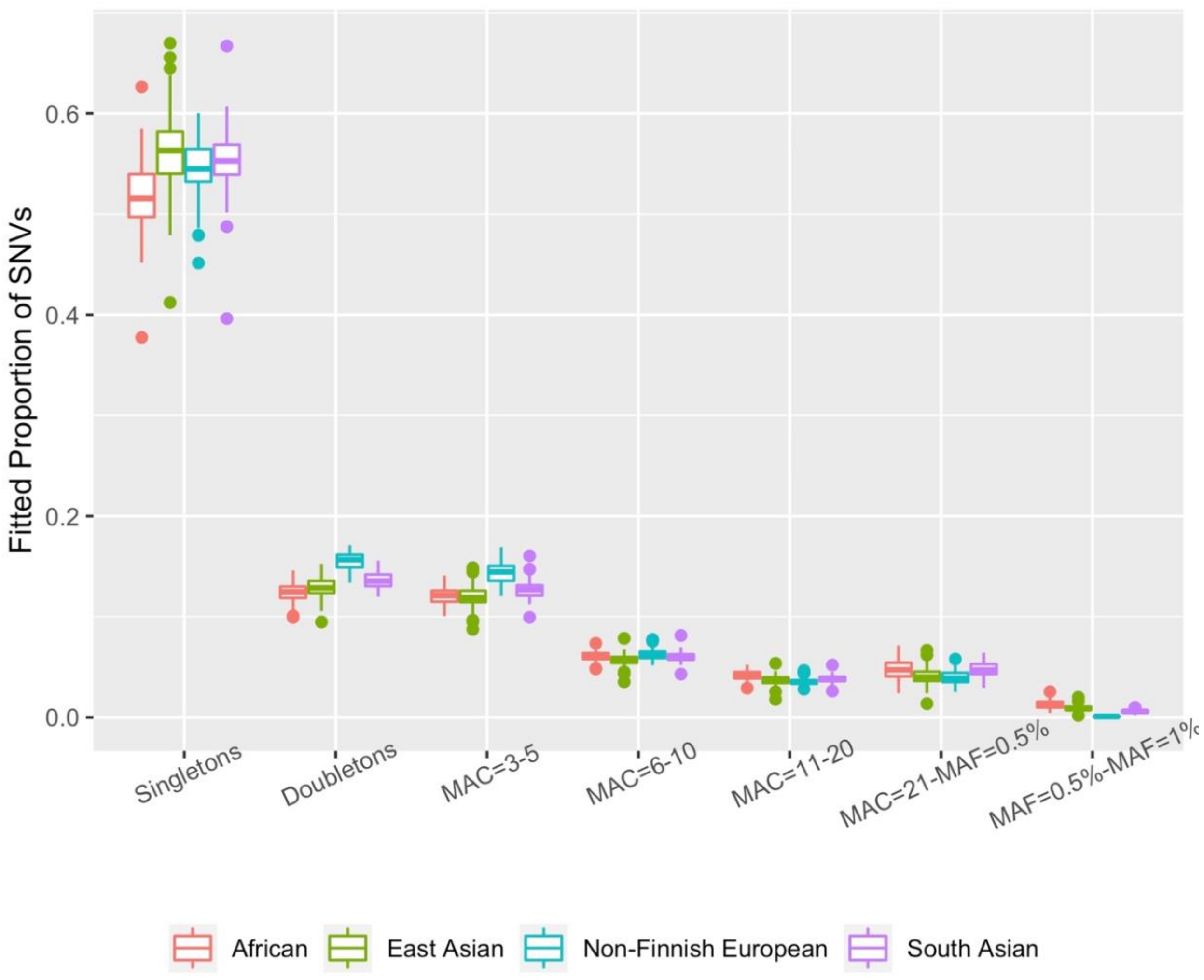


***Supplemental Figure 3. Variation in the AFS estimated proportions*** *The estimated proportion of variants in each MAC bin within the gnomAD target data across ancestry specific blocks are shown. More variation is seen over blocks than ancestries with the most variation seen with the singleton bin. The proportions are similar despite sample size ranging from 8,128 (African) to 56,885 (Non-Finnish European).*

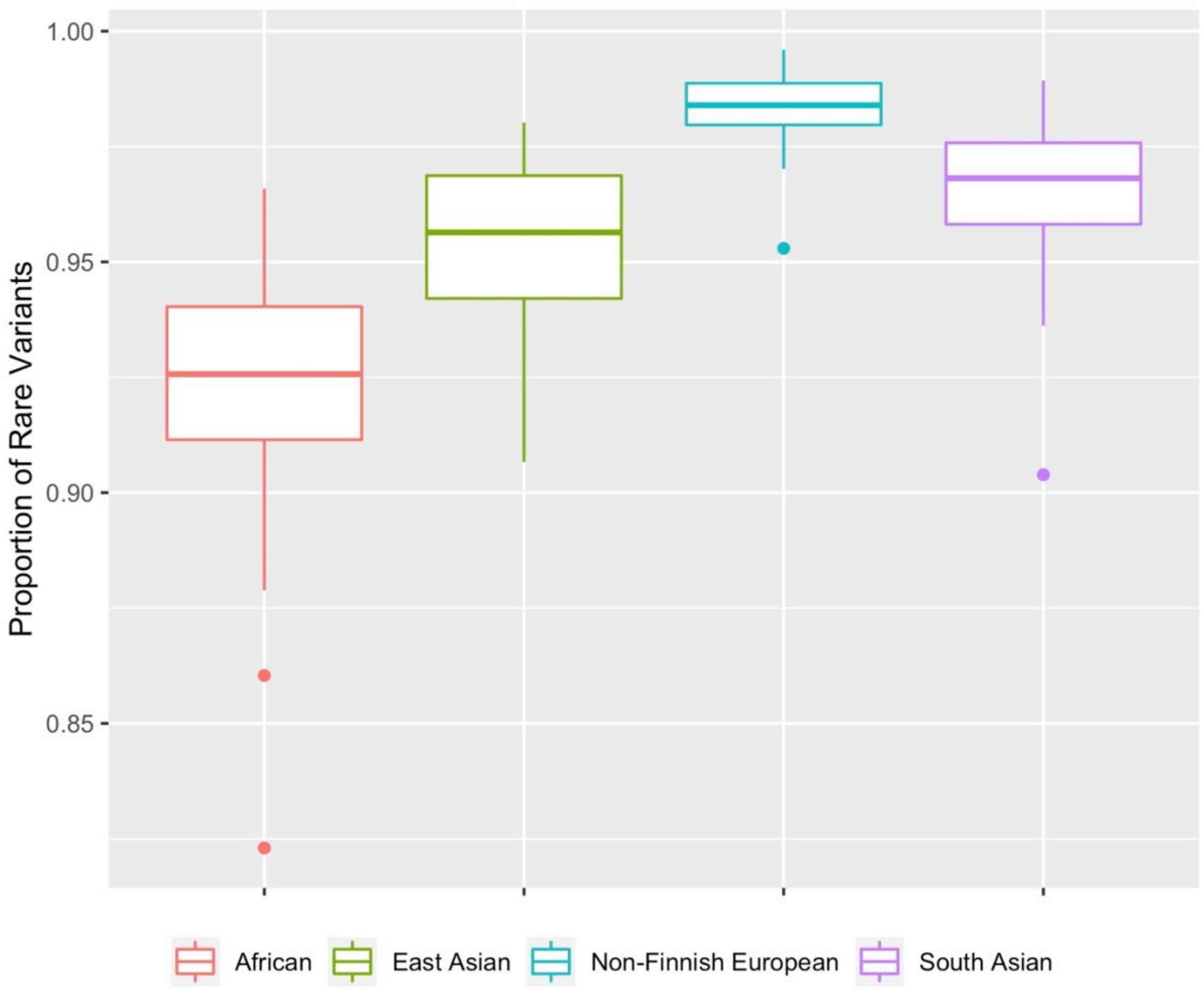


***Supplemental Figure 4. Proportion of Rare Variants*** *The total proportion of rare variants in the gnomAD v2.1 target varies over blocks and between ancestry/sample size groups. Sample sizes also vary: African: N = 8,128; East Asian: N = 9,197; Non-Finnish European: N = 56,885; South Asian: N = 15,308.*

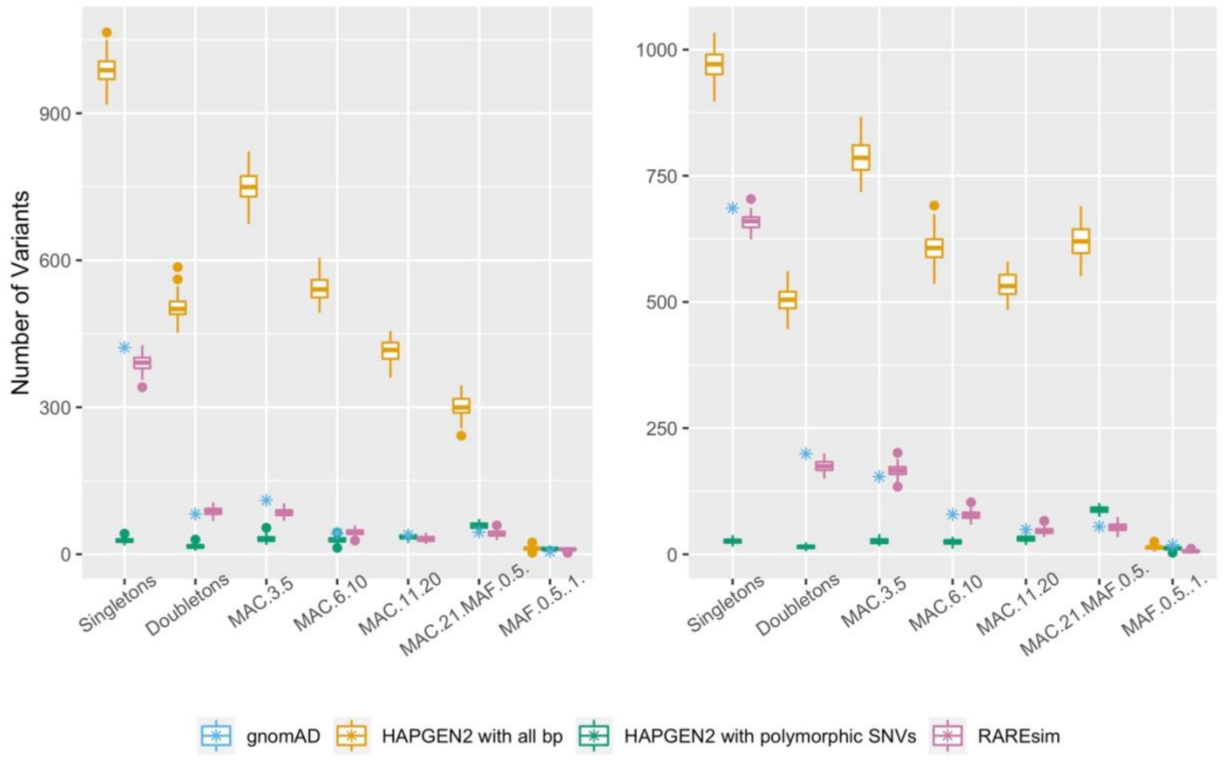


***Supplemental Figure 5. Evaluation of RAREsim*** *For the block with the median number of bp, RAREsim is compared to gnomAD, the default implementation of HAPGEN2 with only polymorphic SNVs, and HAPGEN2 with all sequencing bases. Ancestry specific simulations are shown for East Asian (left) and South Asian (right) ancestry groups, matching the sample size observed in gnomAD to enable a direct comparison of MACs. RAREsim captures the expected number of variants within each MAC bin, while the other simulation methods either grossly underestimate (default implementation of HAPGEN2) or overestimate (HAPGEN2 with all sequencing bp) the number of variants.*

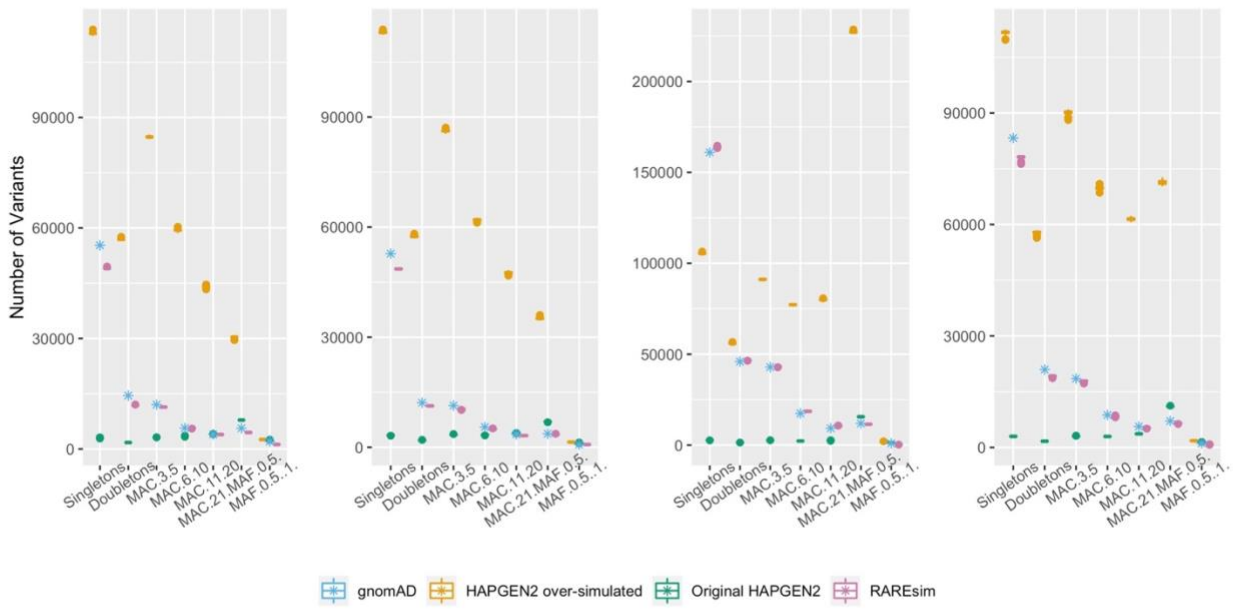


***Supplemental Figure 6. Evaluation of RAREsim across chromosome 19*** *Combining information across all blocks, RAREsim is compared to gnomAD, the default implementation of HAPGEN2 with only polymorphic SNVs, and HAPGEN2 with all sequencing bases. Ancestry specific simulations are shown for (left to right) African, East Asian, Non-Finnish European, and South Asian ancestry groups, matching the sample size observed in gnomAD to enable a direct comparison of MACs. RAREsim captures the expected number of variants within each MAC bin, while the other simulation methods either grossly underestimate (default implementation of HAPGEN2) or overestimate (HAPGEN2 with all sequencing bp) the number of variants.*

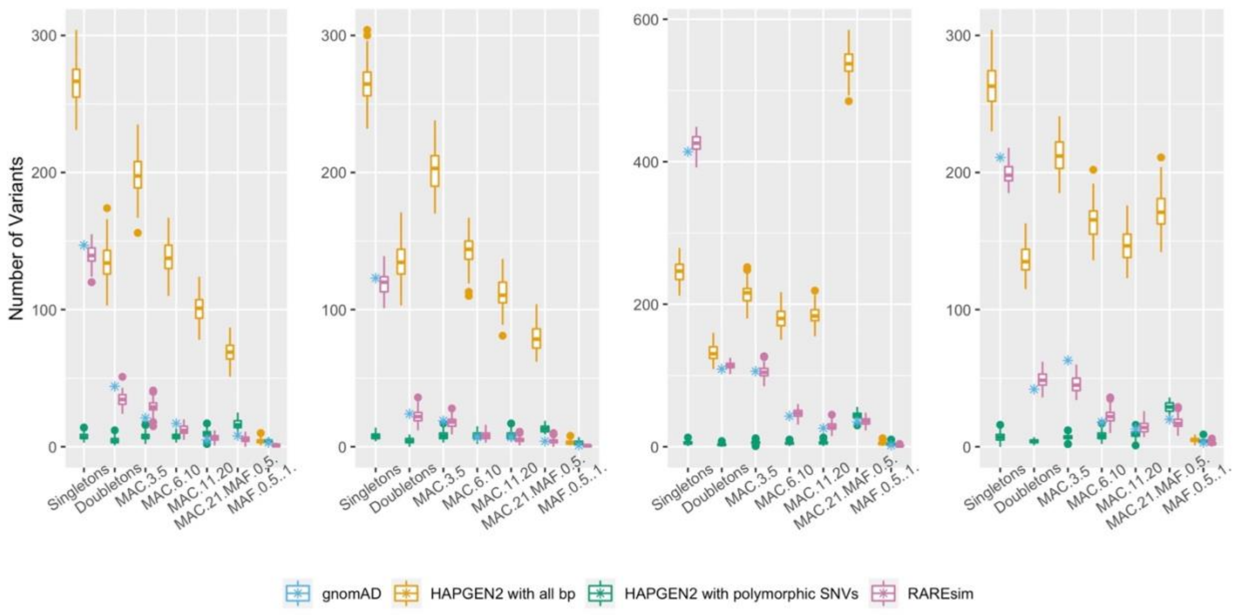


***Supplemental Figure 7. RAREsim evaluation – 5th percentile block*** *Simulations for the block with the 5th percentile in total number of variants is shown here. RAREsim is compared to gnomAD, the default implementation of HAPGEN2 with only polymorphic SNVs, and HAPGEN2 with all sequencing bases. Ancestry specific simulations are shown for (left to right) African, East Asian, Non-Finnish European, and South Asian ancestry groups, matching the sample size observed in gnomAD to enable a direct comparison of MACs. RAREsim captures the expected number of variants within each MAC bin, while the other simulation methods either grossly underestimate (default implementation of HAPGEN2) or overestimate (HAPGEN2 with all sequencing bp) the number of variants.*

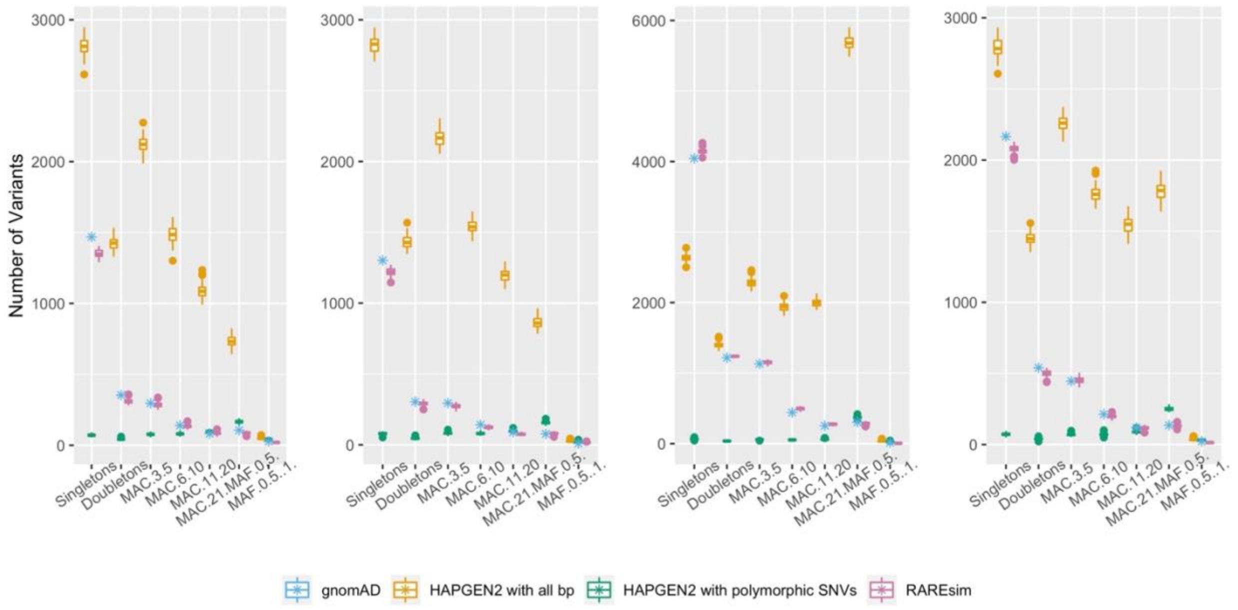


***Supplemental Figure 8. RAREsim evaluation – 95$^{th}$ percentile block*** *Simulations for the block with the 95$^{th}$ percentile in total number of variants is shown here. RAREsim is compared to gnomAD, the default implementation of HAPGEN2 with only polymorphic SNVs, and HAPGEN2 with all sequencing bases. Ancestry specific simulations are shown for (left to right) African, East Asian, Non-Finnish European, and South Asian ancestry groups, matching the sample size observed in gnomAD to enable a direct comparison of MACs. RAREsim captures the expected number of variants within each MAC bin, while the other simulation methods either grossly underestimate (default implementation of HAPGEN2) or overestimate (HAPGEN2 with all sequencing bp) the number of variants.*

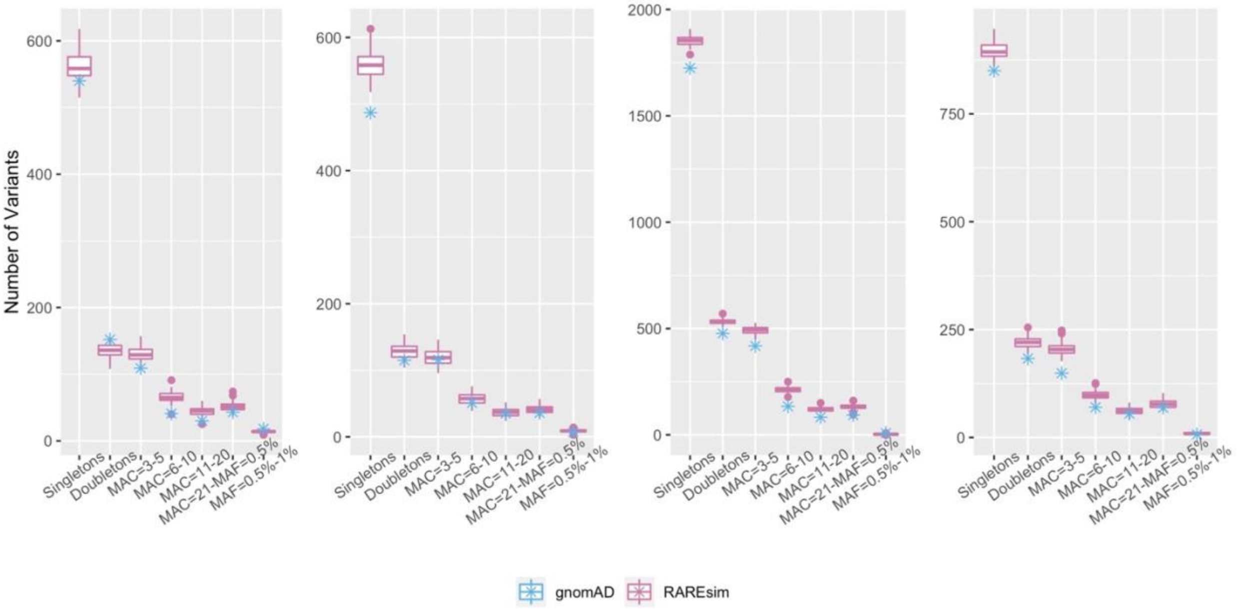


***Supplemental Figure 9. Chromosome 1 simulation with default parameters*** *The coding region of the GENCODE region on chromosome 1 was simulated using ancestry specific default parameters. Ancestry specific simulations are shown for (left to right) African, East Asian, Non-Finnish European, and South Asian ancestry groups, matching the sample size observed in gnomAD to enable a direct comparison of MACs. RAREsim captures the expected number of variants within each MAC bin within the region on chromosome 1 using default parameters derived from chromosome 19.*

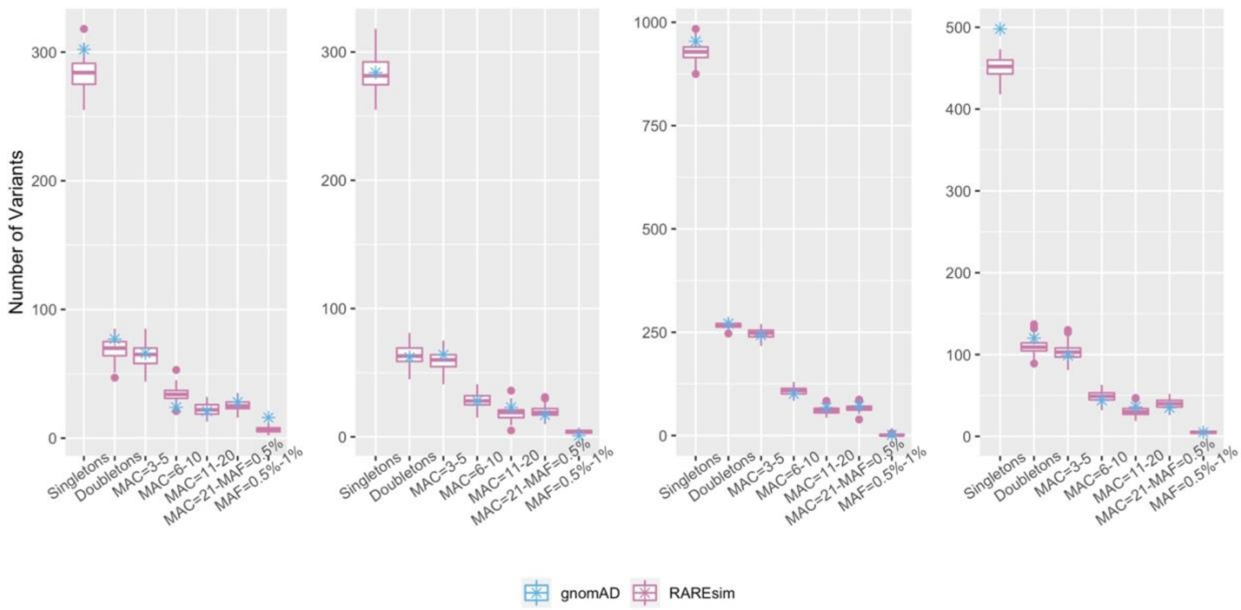


***Supplemental Figure 10. Chromosome 6 simulation with default parameters*** *The coding region of the GENCODE region on chromosome 6 was simulated using ancestry specific default parameters. Ancestry specific simulations are shown for (left to right) African, East Asian, Non-Finnish European, and South Asian ancestry groups, matching the sample size observed in gnomAD to enable a direct comparison of MACs. RAREsim captures the expected number of variants within each MAC bin within the region on chromosome 6 using default parameters derived from chromosome 19.*

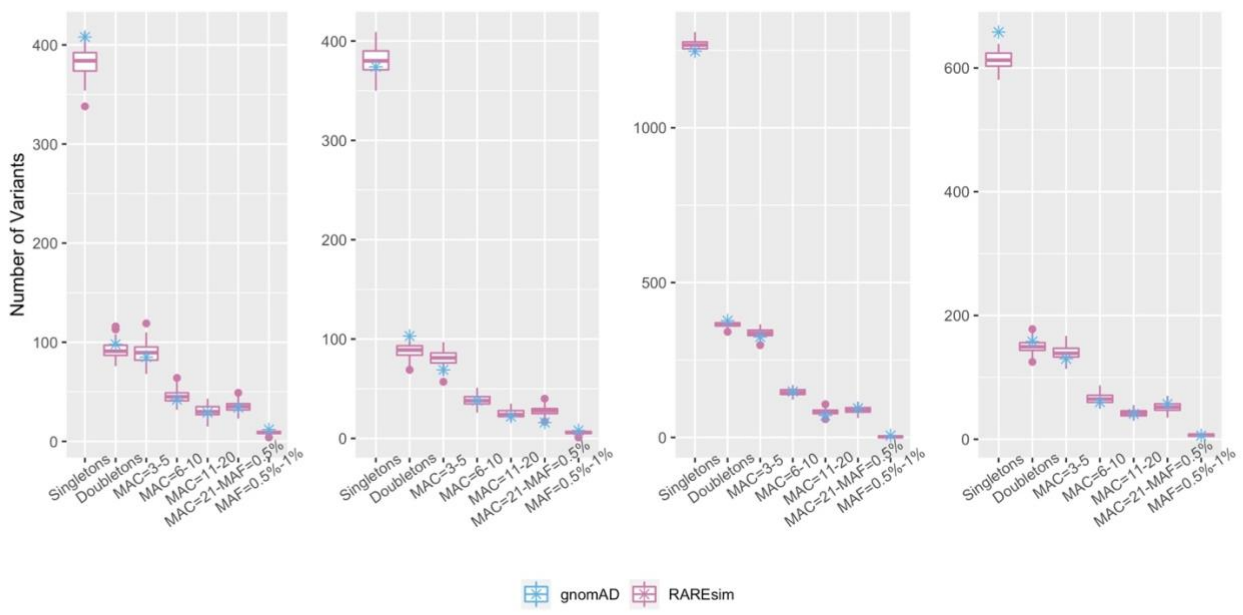


***Supplemental Figure 11. Chromosome 9 simulation with default parameters*** *The coding region of the GENCODE region on chromosome 9 was simulated using ancestry specific default parameters. Ancestry specific simulations are shown for (left to right) African, East Asian, Non-Finnish European, and South Asian ancestry groups, matching the sample size observed in gnomAD to enable a direct comparison of MACs. RAREsim captures the expected number of variants within each MAC bin within the region on chromosome 9 using default parameters derived from chromosome 19.*

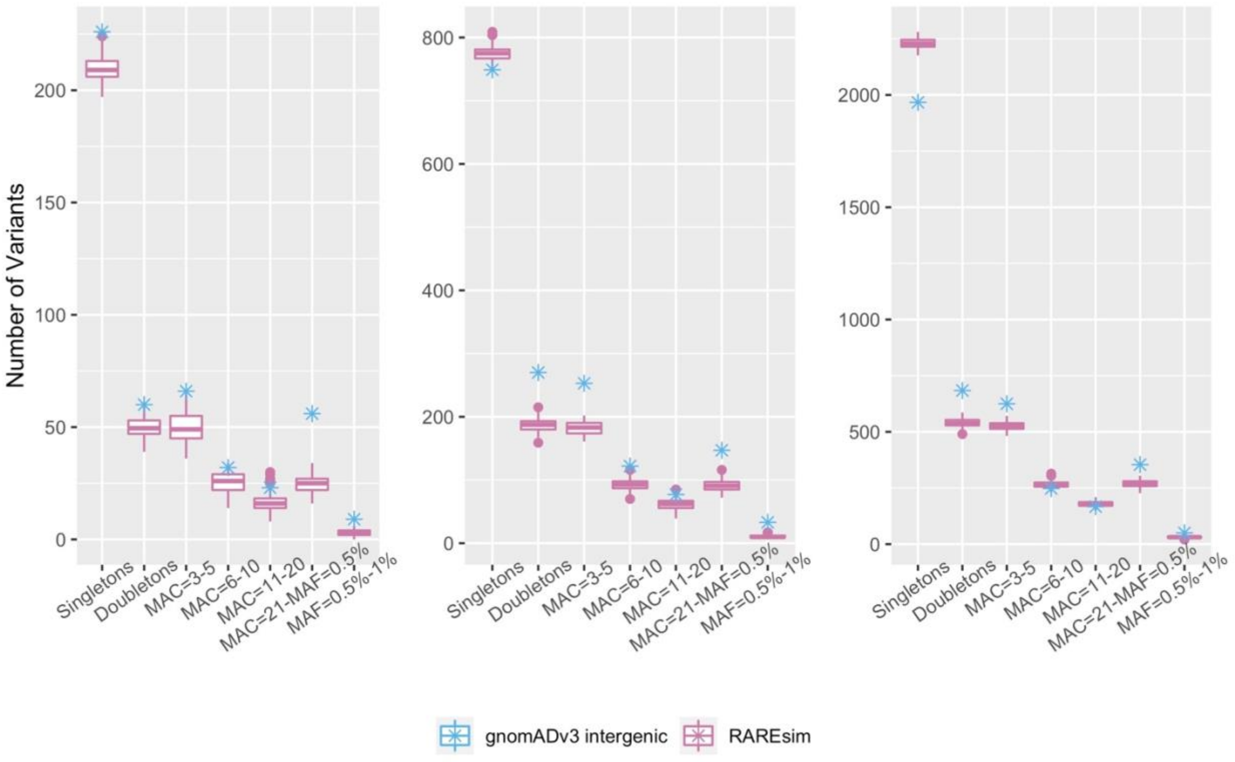


***Supplemental Figure 12. RAREsim default parameters simulating to match an intergenic region in gnomAD v3 African ancestry group*** *Simulations were performed to match the sample size of the gnomAD v3 African ancestry group (N=21,246) for an intergenic region that matched the size of the coding region on that block. The subset of the intergenic portion of the blocks with the 5$^{th}$, median, and 95$^{th}$ percentile of blocks (left to right) were simulated with RAREsim default parameters. RAREsim captures the expected number of variants within each MAC bin for all three blocks.*

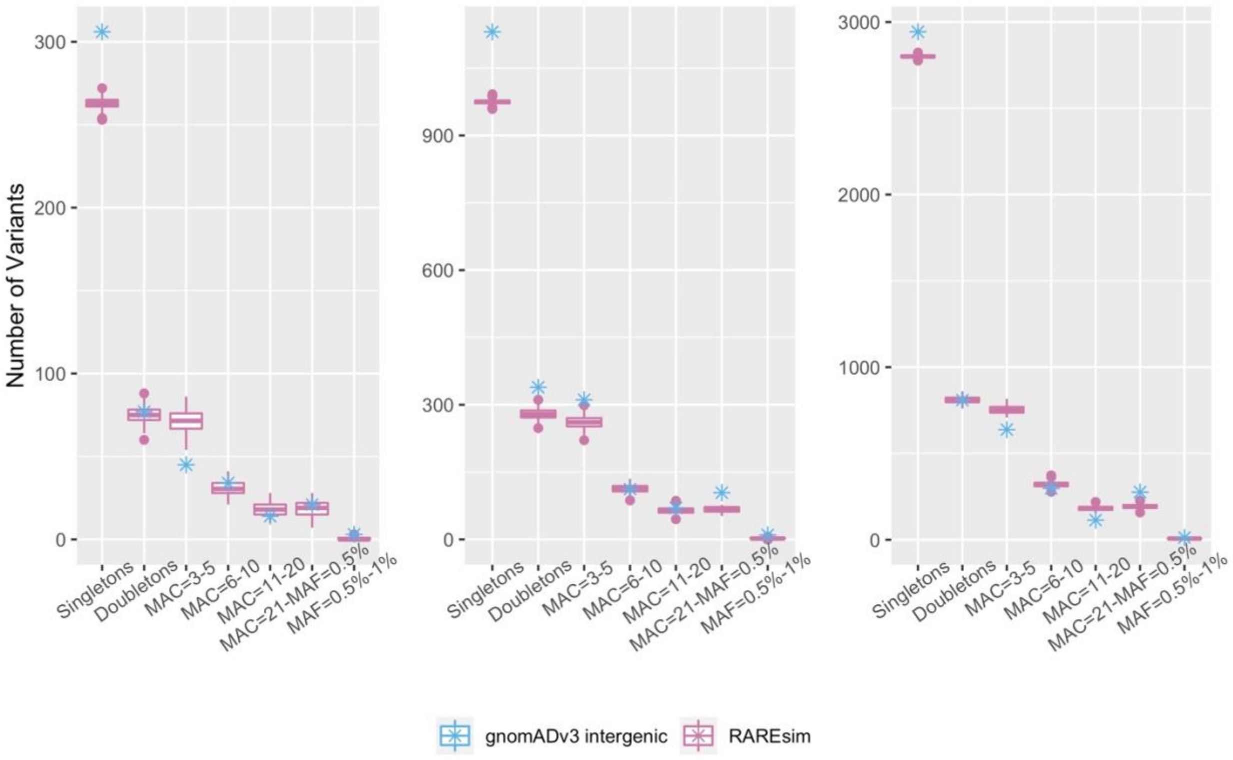


***Supplemental Figure 13. RAREsim default parameters simulating to match an intergenic region in gnomAD v3 Non-Finnish European ancestry group*** *Simulations were performed to match the sample size of the gnomAD v3 Non-Finnish European ancestry group (N=32,299) for an intergenic region that matched the size of the coding region on that block. The subset of the intergenic portion of the blocks with the 5$^{th}$, median, and 95$^{th}$ percentile of blocks (left to right) were simulated with RAREsim default parameters. RAREsim captures the expected number of variants within each MAC bin for all three blocks.*

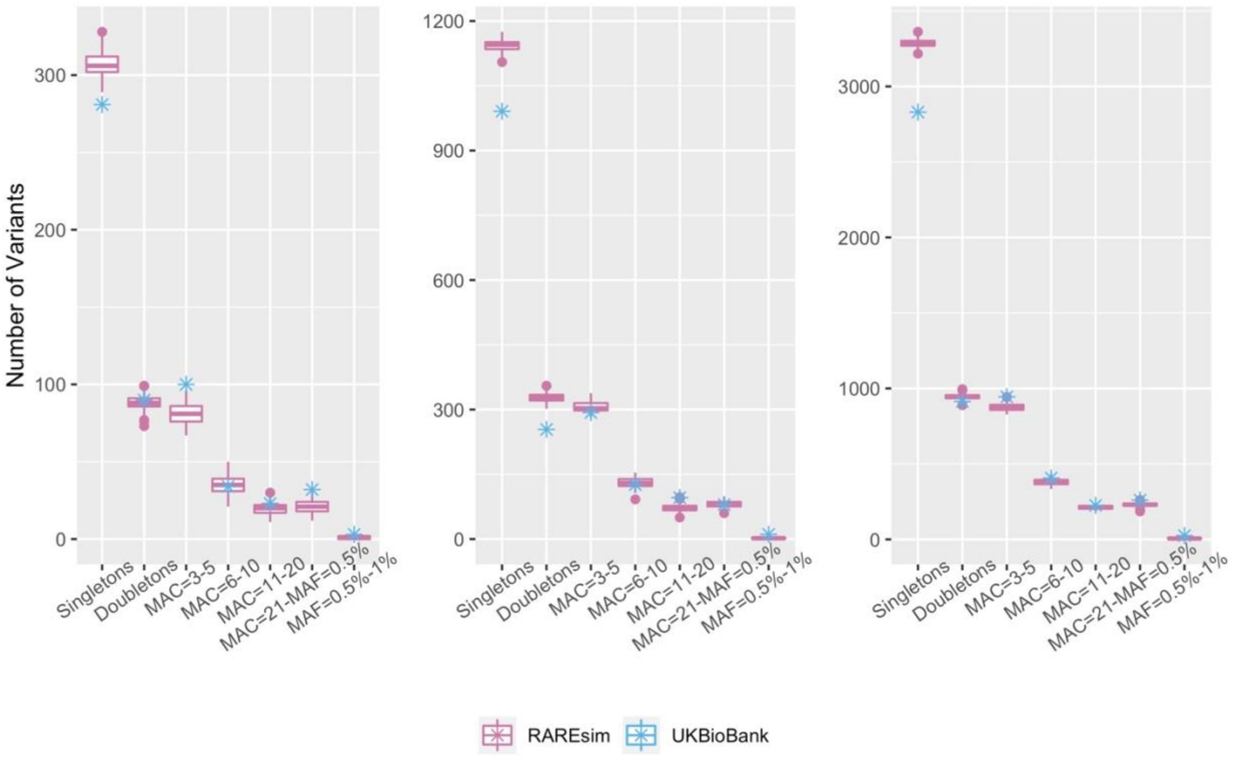


***Supplemental Figure 14. RAREsim default parameters simulating to match UK Biobank data***
*RAREsim was used to simulate a Non-Finnish European sample to match the sample size of the subjects with self-reported British ethnicity (N=41,246). Default parameters were used for the simulation of the block with the 5$^{th}$, median, and 94$^{th}$ percentile number of bp. RAREsim matches the observed UK Biobank data well, with slight overestimations in the number of singletons.*

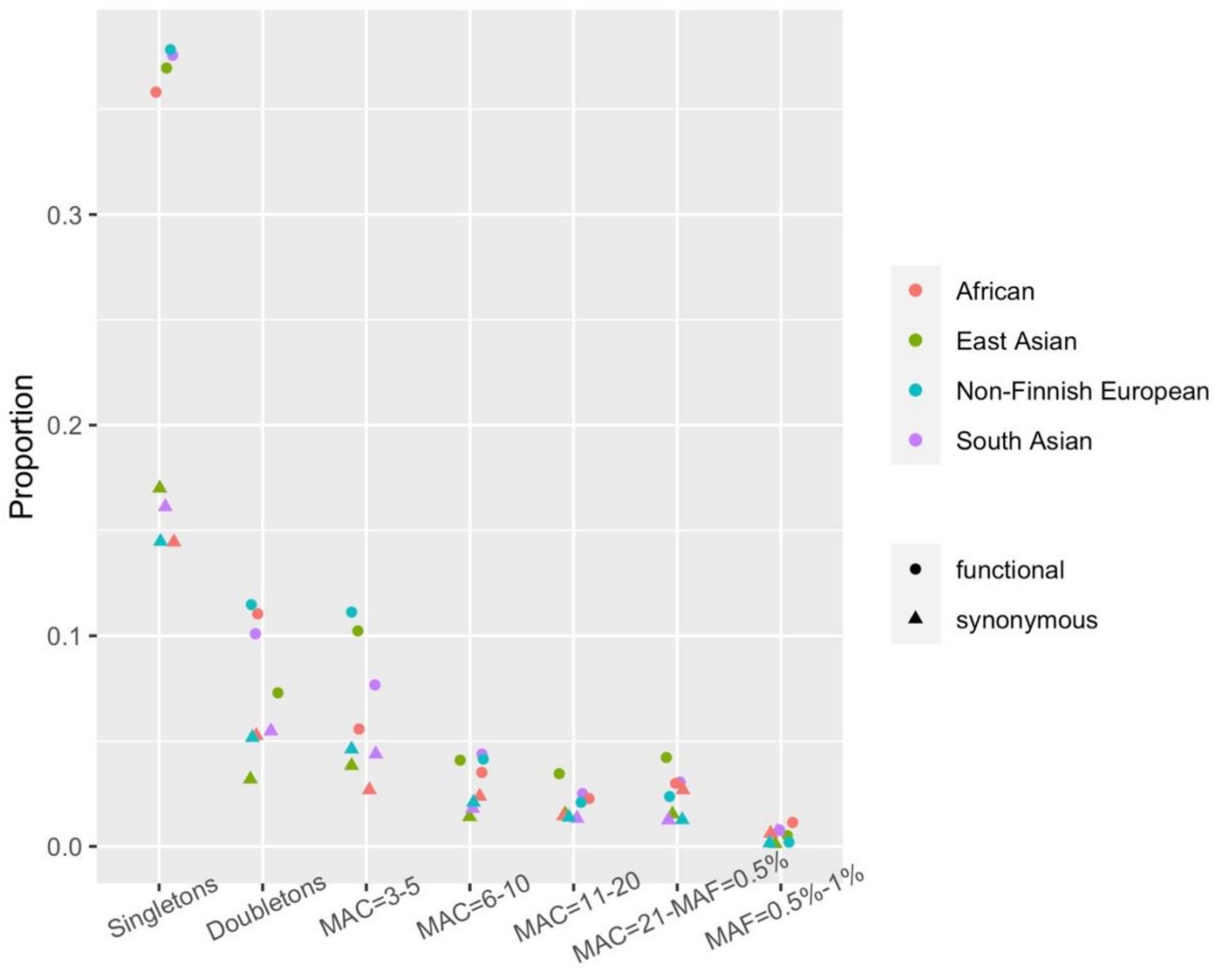


***Supplemental Figure 15. AFS of functional and synonymous variants*** *The proportion of variants in each MAC bin for the block with the median number of bp, separated into functional and synonymous variants and broken down by ancestry.*

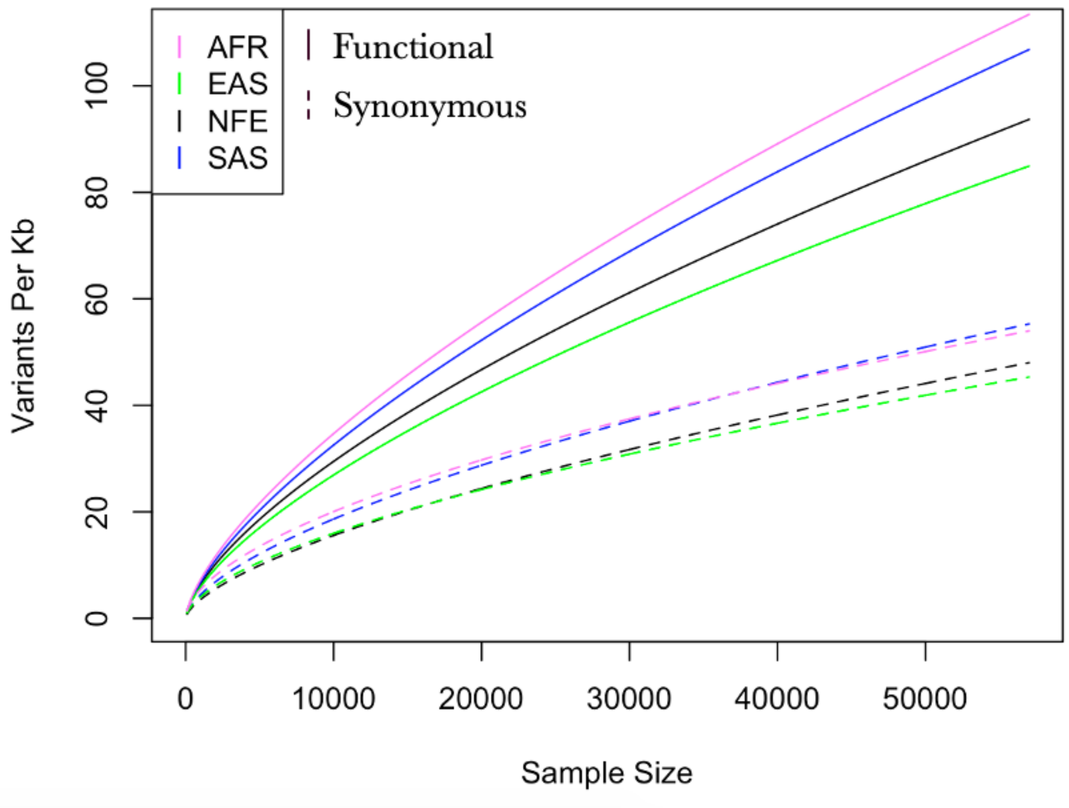


***Supplemental Figure 16. Number of Variants for functional and synonymous variants*** *The Number of Variants functions are shown for the functional (solid line) and synonymous (dotted line) variants for each ancestry for the block with the median number of variants.*

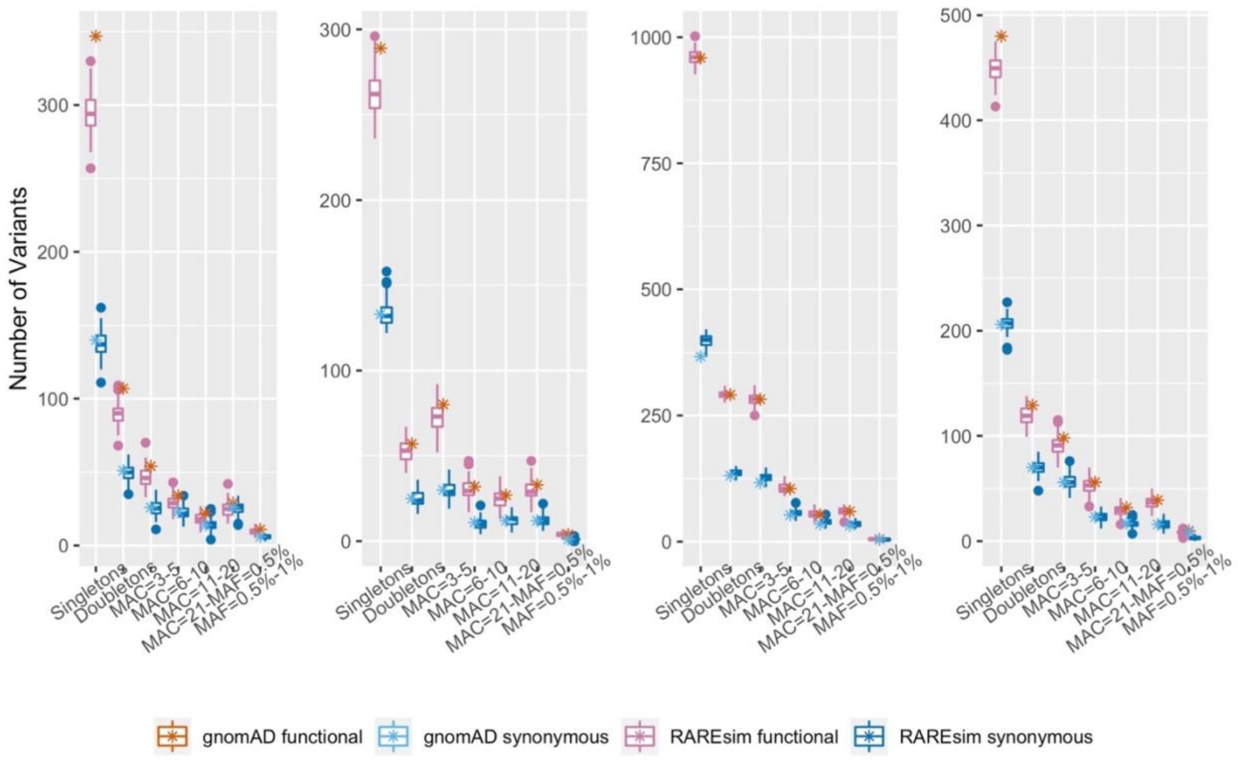


***Supplemental Figure 17. RAREsim simulations stratified over functional and synonymous variants*** *RAREsim was implemented stratifying over functional and synonymous variants. The number of variants in each bin approximates both the number of functional and synonymous variants well across all ancestries, (left to right) African, East Asian, Non-Finnish European, and South Asian.*

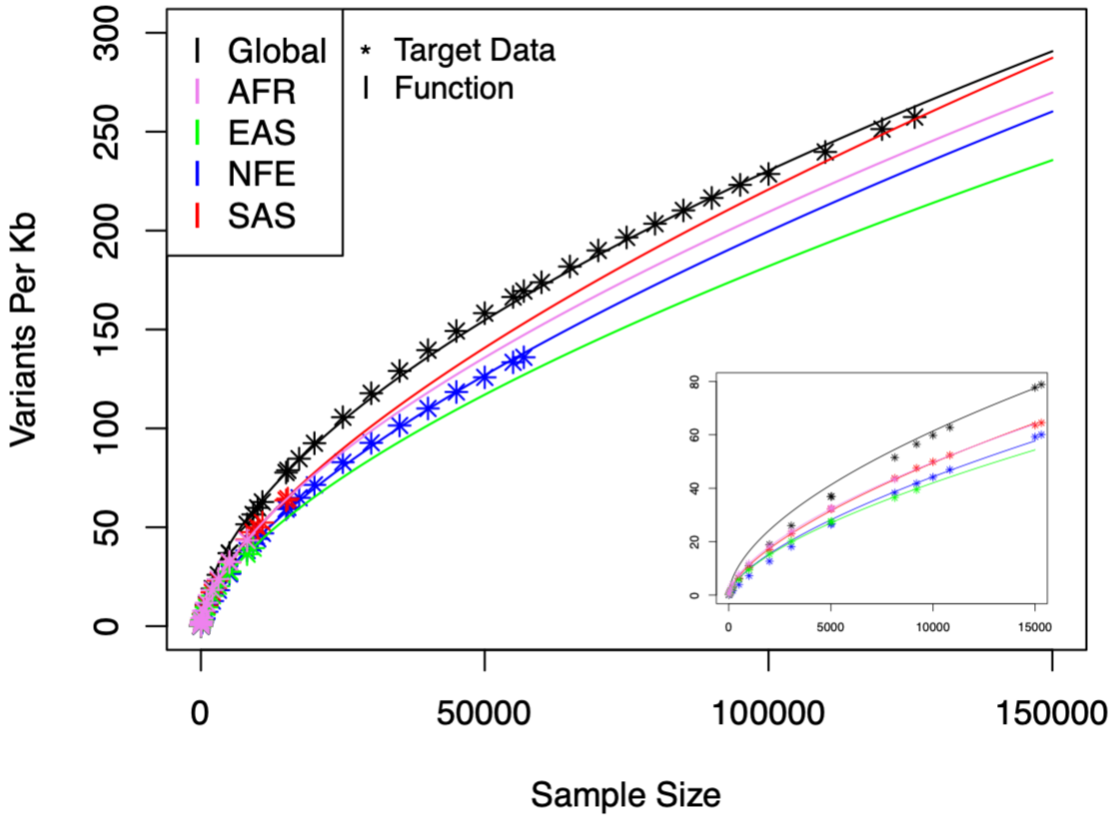


***Supplemental Figure 18. Simulating large sample sizes*** *The Number of Variants function is shown, extrapolating to a sample size of 150,000. The global sample is all gnomAD ancestries combined. The observed median (default) data is shown with stars and the fitted functions with lines. The lower right corner shows the same plot, but with 15,000 samples.*

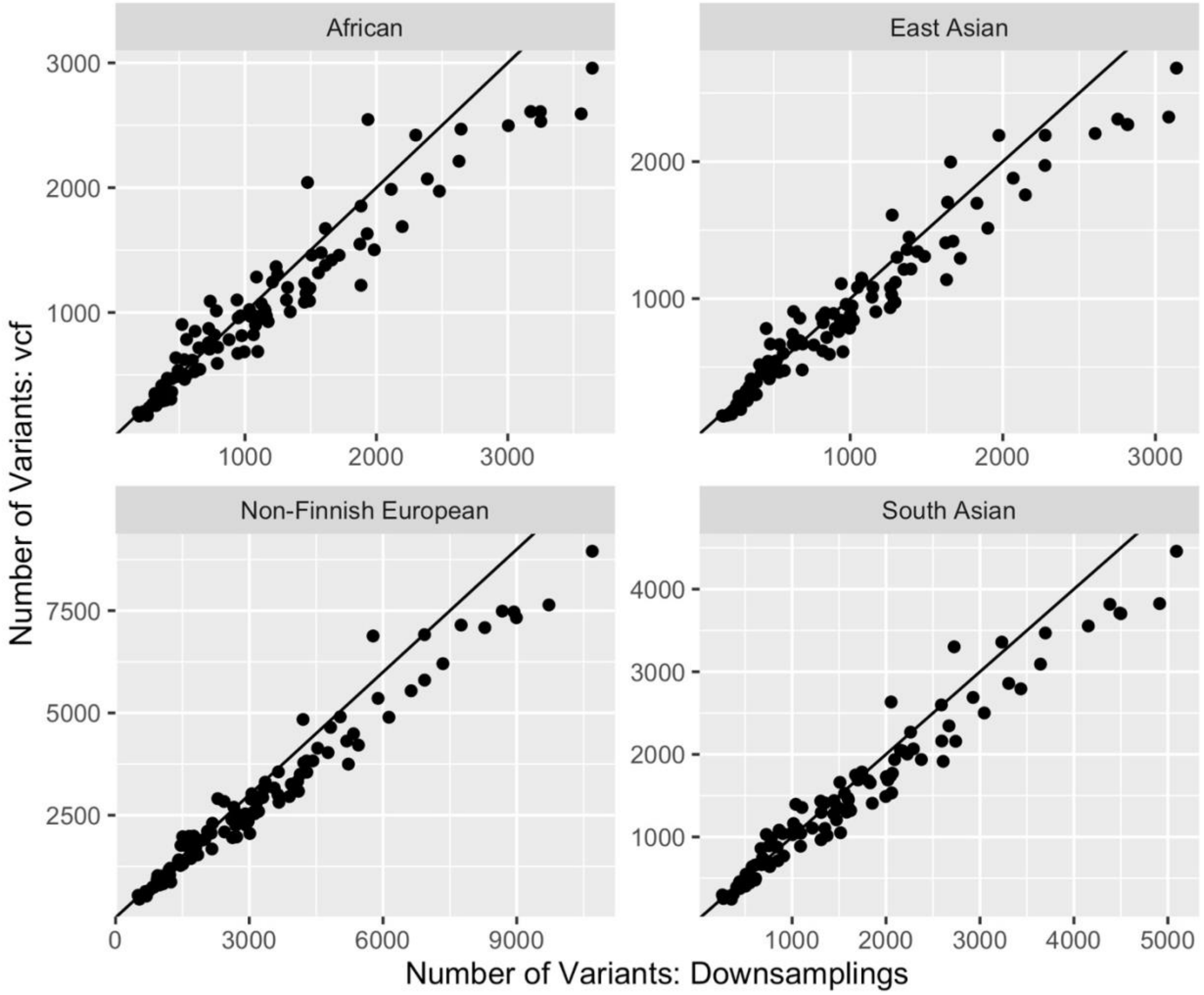


***Supplemental Figure 19. Discrepancy between gnomADv2.1 vcf and downsamplings data***
*The discrepancy in the number of variants between the vcf (y axis) and downsamplings (x axis) data per block from gnomADv2., separated by ancestry.*

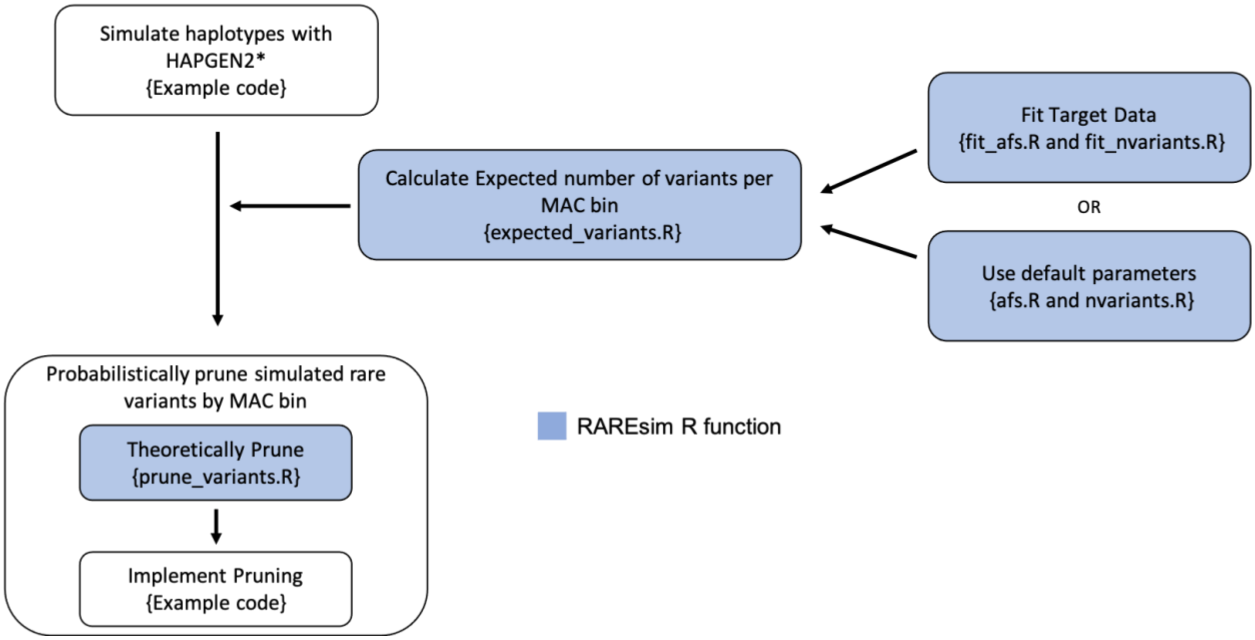


**With all sequencing bases to allow for an abundance of variants*

***Supplemental Figure 20. RAREsim implementation flow chart*** *Flow chart describing the implementation of RAREsim, using the RAREsim R package (blue) and example code on the RAREsim GitHub page (white).*
